# Force dependence of filopodia adhesion: involvement of myosin II and formins

**DOI:** 10.1101/195420

**Authors:** N. O. Alieva, A. K. Efremov, S. Hu, D. Oh, Z. Chen, M. Natarajan, H. T. Ong, A. Jégou, G. Romet-Lemonne, J. T. Groves, M. P. Sheetz, J. Yan, A. D. Bershadsky

**Author notes:** Equal contribution (N. O. Alieva, A. K. Efremov).

## Abstract

Filopodia are dynamic membrane protrusions driven by polymerization of an actin filament core, mediated by formin molecules at the filopodia tips. Filopodia can adhere to the extracellular matrix and experience both external and cell generated pulling forces. The role of such forces in filopodia adhesion is however insufficiently understood. Here, we induced sustained growth of filopodia by applying pulling force to their tips via attached fibronectin-coated beads trapped by optical tweezers. Strikingly, pharmacological inhibition or knockdown of myosin IIA, which localized to the base of filopodia, resulted in weakening of filopodia adherence strength. Inhibition of formins, which caused detachment of actin filaments from formin molecules, produced similar effect. Thus, myosin IIA-generated centripetal force transmitted to the filopodia tips through interactions between formins and actin filaments are required for filopodia adhesion. Force-dependent adhesion led to preferential attachment of filopodia to rigid versus fluid substrates, which may underlie cell orientation and polarization.

## INTRODUCTION

Filopodia are ubiquitous cell extensions involved in cell motility, exploration of the microenvironment and adhesion ^1, 2^. These finger-like membrane protrusions help cells to determine the direction of movement ^3^, establish contacts with other cells ^4, 5^ and capture inert particles or living objects (bacteria), which cells subsequently engulf ^6–9^. Filopodia are involved in numerous processes of embryonic development, as well as in cell migration in adult organisms. Moreover, augmented filopodia activity is a hallmark of tumor cells, which use them in the processes of invasion and metastasis^1^.

The main element of filopodia is the actin core, which consists of parallel actin filaments with barbed ends oriented towards the filopodium tip, and pointed ends toward the cell body ^1, 2, 10^. Actin filaments are connected to each other by several types of crosslinking proteins ^11–14^ The filopodia grow via actin polymerization at the tip, in a process driven by formin family proteins such as mDia2 ^15–17, 18^, FMNL2 & 3 ^19–21^, as well as by actin elongation protein Ena/VASP ^15, 22, 23, 24^. In addition to proteins that crosslink and polymerize actin, filopodia also contain actin based molecular motors, such as myosin X, localized to the tips of the filopodia ^25^. Although the function of myosin X is unclear, it is known to be required for filopodia growth, and its overexpression promotes filopodia formation ^26, 27^.

Adhesion of the filopodia to the extracellular matrix (ECM) is mediated by the integrin family of receptors (e.g. α_v_β_3_) ^25, 28^, which are localized to the tip area. One possible function of myosin X is the delivery of integrins to this location ^25^. In addition to integrins, filopodia tips have been shown to contain other proteins involved in integrin mediated adhesion, such as talin ^29^ and RIAM ^30^. Several studies suggest that typical cell matrix adhesions, known as focal adhesions, could in some cases originate from filopodia ^31, 32^. Thus, filopodia could be considered as primary minimal cell matrix adhesion structures.

The hallmark of integrin mediated adhesions of focal adhesion type is their mechanosensitivity ^33–35^. They grow in response to pulling forces applied to them, either by the actomyosin cytoskeleton, or exogenously by micromanipulations, and may play a role in matrix rigidity sensing. Indeed, correlation between focal adhesion size and matrix rigidity is well-documented ^36–38^. Filopodia also may participate in matrix rigidity sensing. For example, it was demonstrated that cell durotaxis, a preferential cell movement along a gradient of substrate rigidity is mediated by filopodia ^39^. However, force dependence of filopodia adhesion has not yet been explored.

In the present study, we monitored filopodia adhesion and growth under conditions of pulling with a constant rate. We have demonstrated that adhesion of filopodia to the ECM strongly depends on myosin II activity and found myosin II filaments localized to the base regions of filopodia. Moreover, formin family protein activity at the filopodia tips is also required for filopodia adhesions, most probably through a role in the transmission of force through the actin core, from the filopodium base to the filopodium tip. Thus, filopodia are elementary units demonstrating adhesion-dependent mechanosensitivity.

## RESULTS

### Dynamics of filopodia induced by expression of myosin X in HeLa-JW cells

Transfection of HeLa-JW cells with either GFP-myosin X or mApple-myosin X resulted in a strong enhancement of filopodia formation in agreement with previous studies ^40^. During filopodia movement, myosin X was concentrated at the filopodia tips, forming characteristic patches sometimes also called “puncta” or “comet tails” (fig. S1A, movie S1). Here, we focused on filopodia originating from stable cell edges and extending along the fibronectin-coated substrate. These filopodia demonstrated periods of persistent growth, with an average velocity of 67 ± 6 nm/s (mean ± SEM, n = 89) interrupted by pauses and periods of shrinking with an average velocity of 28 ± 3 nm/s (mean ± SEM, n = 100). This behavior is consistent with previously published results ^41^. In addition to myosin X, the filopodia tips were also enriched in several other proteins such as mDia2, VASP and talin (fig. S1B and D).

To observe the dynamics of filopodia adhesion and protrusion under controlled experimental conditions, we monitored the growth of filopodia that were adhered to fibronectin-coated beads trapped by optical tweezers (see supplementary information for more details). First, 2μm diameter fibronectin-coated polystyrene beads were placed onto filopodia tips by the optical tweezers. After 20-30 s, which is required for the initial attachment of the bead to the filopodium, the movement of microscope piezo stage in the direction from the tip to base of filopodium was initiated (Fig. 1, movies S2, S3, S9A). The force exerted by filopodium on the bead was monitored by measuring the bead displacement from the center of the trap (Δx). In order to preserve the structural integrity of the filopodia, the velocity of the stage movement was set to approximately 10-20nm/s, which is slower than the average velocity of spontaneous filopodia growth. With this setup we observed sustained filopodia growth for more than 10 mins, during which time the tdTomato-Ftractin labelled actin core remained intact (Fig. 1B and 3A, movie S2). Pulling-induced filopodia growth was depended on integrin-mediated adhesion of filopodia tips to fibronectin-coated beads. When the beads were coated with concanavalin A instead of fibronectin, application of force never induced the growth of filopodia actin cores. Instead, pulling via concanavalin A-coated bead resulted either in detachment of filopodia tips from the beads (18%), or withdrawal of the bead from the trap (27%), or, in majority of cases (55%), in the formation of membrane tethers, n = 11 (movie S4).

**Fig. 1.**
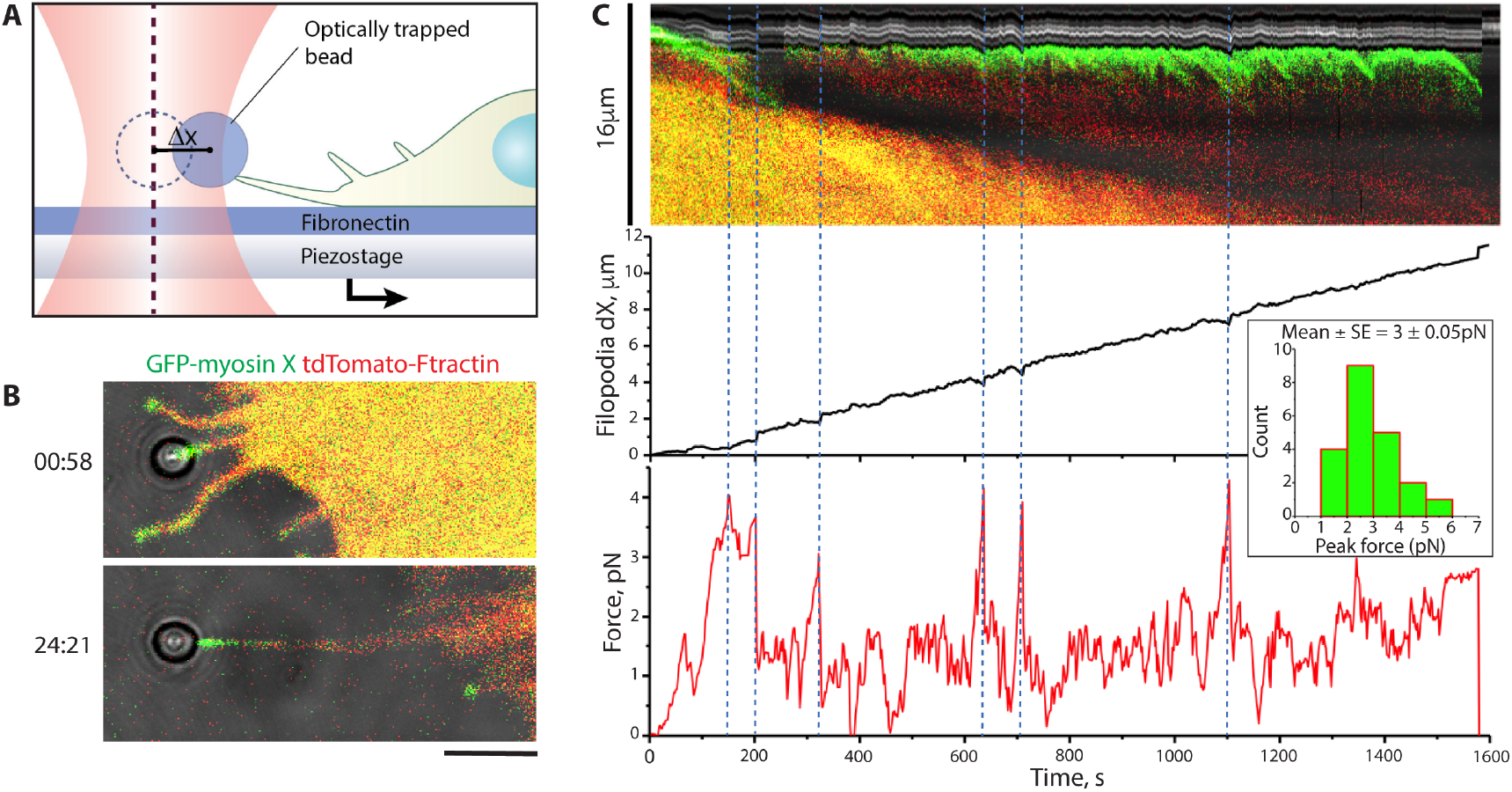
Dynamics of pulling-induced filopodia growth. **(A)** Experimental setup used to observe force-induced filopodia growth. Optical tweezers were used to trap fibronectin-coated microbeads attached to filopodia tips of HeLa-JW cells. **(B)** Confocal images of a typical cell expressing GFP-myosin X and tdTomato-Ftractin with an attached bead, taken immediately after starting of stage movement (top) and in the course of sustained growth (bottom). Note that both myosin X and actin remain at the filopodium tip during growth. See also movies S2-3. Scale bar, 5μm. **(C)** Top panel: A kymograph showing the dynamics of myosin X and actin in the filopodium shown in (B). This kymograph is composed from two parts smoothly combined next to each other: line was drawn through basis region (left part), and - proximal region of the filopodium (right). Middle panel: Filopodium growth in relation to the coordinate system of the microscope stage. The origin of the coordinate system corresponds to the bead position in the center of the laser trap at the initial time point. The coordinate of the bead is changing due to the uniform movement of the stage, and fluctuations of the bead position inside the trap. Lower panel: Forces experienced by the bead. Note the discrete peak force values corresponding to the moments of filopodia growth cessation (seen in the middle panel) as marked with dotted lines. Inset: The distribution of peak force values, based on the pooled measurements of 21 peaks from 6 beads.

**Fig. 2.**
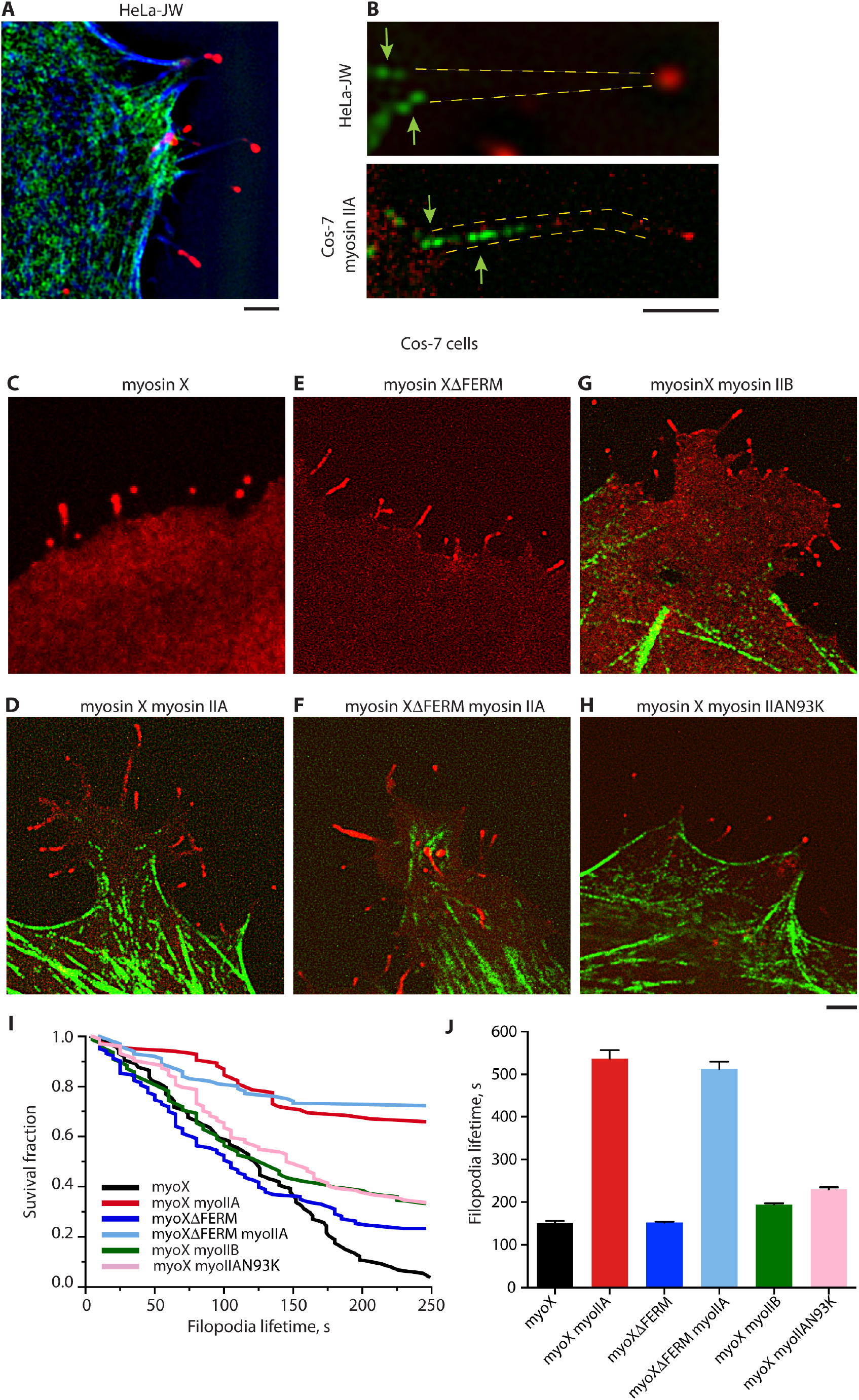
Myosin II filaments in filopodia of HeLa-JW and Cos-7 cells. **(A)** Visualization of mApple-myosin X (shown in red), RLC-GFP (green) and mTagBFP-Lifeact (blue) in HeLa-JW cell. **(B)** Zoomed images of bipolar myosin IIA filaments at the bases of filopodia. Upper panel: myosin IIA and myosin X were labeled as indicated above in HeLa-JW cell. Lower panel: Cos-7 cell expressing mApple-myosin X (red) and GFP-myosin IIA heavy chain (green). Arrows indicate myosin II mini-filaments. **(C)-(H)** Images of Cos-7 cells expressing myosin X (red) and myosin IIA or IIB (green) and their mutants: **(C)** GFP-myosin X. **(D)** mApple-myosin X and GFP-myosin IIA heavy chain. **(E)** GFP-myosin X ΔFERM. **(F)** GFP-myosin X ΔFERM and mCherry-myosin IIA heavy chain. **(G)** mApple-myosin X and GFP-myosin IIB heavy chain. **(H)** mApple-myosin X and GFP-myosin IIA N93K. Note the presence of myosin IIA filaments for A, B, D, F and H and the absence of myosin IIB at the filopodia bases. See also movies S8A-G, which correspond to images A, C-H, respectively. Scale bars, 2μm. **(I)** Survival fraction of the filopodia cohort in Cos-7 cells with the above constructs. **(J)** Lifetimes of filopodia in cells transfected with different constructs of myosin II and myosin X. To calculate the average filopodia lifetime, experimentally measured survival fraction of cell filopodia was fitted to the exponential decay function: survival fraction = *e*^−t/λ^, where t is time and λ is the average filopodia lifetime. Calculated lifetime values are (mean±SD) 151.0±5.1 (GFP-myosin X only, n=72, 3 cells), 537.2±19.8 (n=30, 3 cells), 152.4±1.7 (n=81, 3 cells), 513.0±16.9 (n=29, 3 cells), 194.7±2.6 (n=53, 3 cells), and 229.8±4.7 (n=70, 4 cells) seconds (in order corresponding to images C-H). All images and data for analysis were collected using structural illumination microscopy (SIM) except (C), which was obtained by spinning disk confocal microscopy (SDCM).

**Fig. 3.**
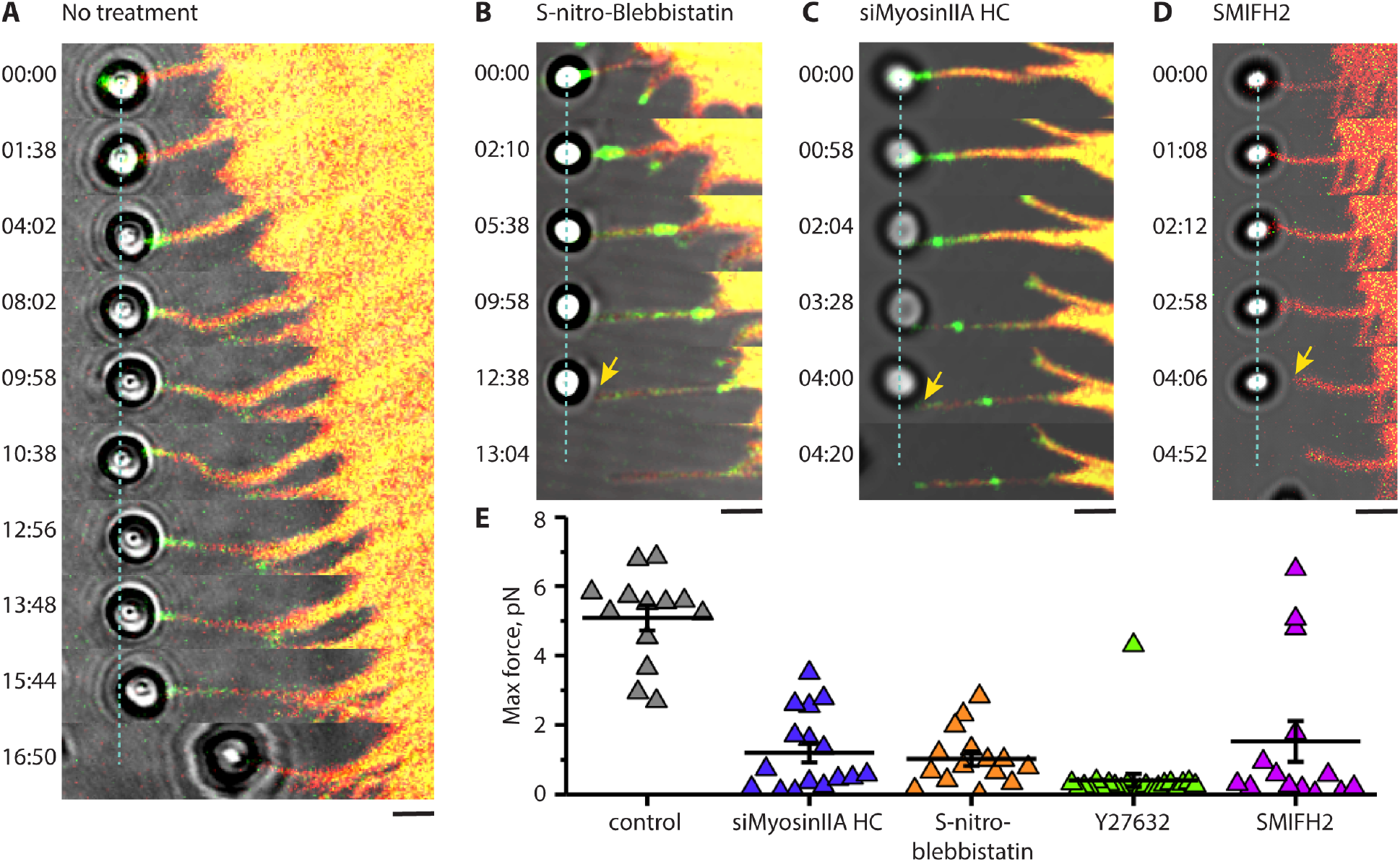
Inhibition of myosin II or formin reduces filopodia adhesion. **(A)** Filopodium growth upon application of pulling force in HeLa-JW cells. The deflection of the bead from its initial position at the center of the laser trap (dashed line) is proportional to the forces exerted by the filopodium (see Fig. 1, movie S6A). At 16:50 min the filopodium retracted and pulled the bead out of the trap. **(B-D)** Filopodia in cells with suppressed myosin II or formin activity cannot maintain sustained adhesion to the bead and do not produce forces sufficient for noticeable bead deflection during the stage movement. Cells treated with 20μM of S-nitro-blebbistatin for 10-20 min (B), transfected with myosin IIA siRNA (C), or treated for with 40μM of the formin inhibitor SMIFH2 for 1 hour (D) are shown. GFP-myosin X and tdTomato-Ftractin are shown in green and red, respectively. See also movies S6B-D. The adhesions of the beads to filopodia were broken at 6 min 45 s, 3 min 50s and 2 min after starting the stage movement for S-nitro-blebbistatin-treated, myosin IIA knocked down, and SMIFH2-treated cells, respectively. Scale bars, 2μm. **(E)** Peak values of the forces exerted by filopodia on the beads during the stage movement in control cells (no treatment) and in cells transfected with myosin IIA siRNA, or treated with S-nitro-blebbistatin, Y27632 (30μM, 10-20 min), or SMIFH2. Mean values (horizontal lines) and SEMs (error bars) are indicated. The mean±SEM of the maximal forces exerted by control filopodia (5.1±0.4pN, n = 13) was significantly higher than those in myosin IIA knockdown, as well as S-nitro-blebbistatin-, Y27632-, and SMIFH2-treated cells (1.2±0.3, n = 16; 1.0±0.2, n = 15; 0.4±0.2, n = 22; and 1.5±0.6pN, n = 14, respectively) (p<0.0001 for control vs all treatment cases).

During the first 3 mins after stage movement commenced, the exerted force approached the maximal value of 3-5 pN. However, it then dropped to the 1.5-2pN range, and remained at this level for a further 1-3 mins, after which it rapidly increased again (Fig. 1C). In a typical experiment, we detected 2-4 such peaks with a mean peak force value of 3pN alternating with the 1-3 min periods of lower force (1.5-2pN).

The pattern of force dependent filopodia elongation described above was typical for myosin X-induced filopodia, but not for filopodia induced by constitutively active Cdc42 (Q61L). Cdc42-induced filopodia attached to laser-trapped fibronectin-coated beads did not grow upon stage movement and eventually pulled the beads out of the trap (movie S5). In this study, we exclusively focused on the filopodia-induced by myosin X overexpression.

Immediately after attachment of the bead to the filopodium tip, the myosin X patch, or a significant portion that pinched off the main myosin X mass, started to move centripetally with an approximate velocity of 31 ± 5 nm/s (mean ± SEM, n = 42). Co-expression of myosin X with VASP in HeLa-JW cells revealed that the retrogradly moving myosin X patches were always colocalized with the patches of VASP (fig. S1B middle panel). Moreover, centripetal movement of myosin X patches in cells transfected with photoactivatable β-actin and myosin X proceeded with the same velocity as the movement of a photoactivated actin spots in the same filopodia (fig. S1C, movie S7). However, despite the retrograde movement of a large part of myosin X, it did not entirely disappear from the filopodium tip and the original amount was fully restored after several minutes (Fig. 1C, kymograph), even though detachment and subsequent centripetal movement of myosin X portions from the filopodium tip were occasionally observed throughout the entire period of force-induced filopodium growth (movie S3).

### Involvement of myosin II in filopodia dynamics

Expression of GFP labeled myosin light chain in HeLa-JW cells showed that myosin II does not localize to the filopodia tips or shafts, but is often located at the proximal ends of the filopodia (Fig. 2A and B top, movie S8A). Structure illumination microscopy (SIM) revealed few myosin II mini-filaments on either side of the filopodium base.

Localization of exogenous myosin IIA at the filopodia bases was especially prominent in Cos-7 cell model. In should be noted that Cos-7 cells express myosin IIB and IIC, but do not contain myosin IIA ^42, 43^. Upon transfection with myosin X, the wild type Cos-7 cells form filopodia with a short life-time that apparently do not adhere to the fibronectin-coated substrate (Fig. 2C, movie S8B). Co-transfection of these cells with GFP-myosin IIA heavy chain resulted in formation of numerous bipolar myosin IIA filaments frequently localized to the bases of filopodia (Fig. 2B bottom, 2D, movie S8C). Majority of filopodia in such cells were associated with myosin IIA filaments during the period of observation. Indeed, the analysis of the histories of 25 filopodia from the movie S8C revealed that 17 of them (68%) had one or more myosin IIA doublets at their bases during the movie duration (4 min 5 s). Myosin X-containing filopodia in Cos-7 cells co-transfected with myosin IIA were somewhat longer than those in wild type Cos-7 lacking myosin IIA (fig. S2). Moreover, expression of myosin IIA significantly increased the lifetime of myosin X-induced filopodia (Fig. 2J) as it can be inferred from the analysis of survival fraction of the filopodia cohort (Fig. 2I).

In agreement with previous studies, formation of filopodia in Cos-7 cells can be also induced by expression of truncated myosin X lacking FERM domain (Fig. 2E, movie S8D) ^44^ Co-expression of myosin IIA significantly enhanced the lifetime of the filopodia induced by this truncated myosin X (Fig. 2I and J, movie S8E) showing that binding of myosin X to integrin via FERM domain ^25^ is not required for myosin IIA-driven enhancement of filopodia lifetime. In contrast, expression of GFP-myosin IIB heavy chain in Cos-7 cells did not affect the filopodia lifetime (Fig 2I, J). Consistently, myosin IIB was not localized to the filopodia base in these cells (Fig. 2G, movie S8F).

Mutant myosin IIA N93K has reduced myosin ATPase and motor activity, but preserves the ability to bind to actin filaments ^45, 46^. GFP fusion construct of this mutant forms filaments localized to the bases of myosin X-induced filopodia in Cos-7 cells, similarly to wild type myosin IIA (Fig. 2H). However, unlike the wild type, the mutant myosin IIA N93K only slightly enhanced the lifetime of filopodia (Fig. 2I, J). Thus, motor activity of myosin IIA is critically important for the effect of myosin IIA on filopodia life time.

We further studied how the presence and activity of myosin IIA affects unconstrained and force-induced filopodia growth. The function of myosin II was suppressed in HeLa-JW cells in three separate experiments: through the inhibition of ROCK by Y27632, by siRNA mediated knockdown of myosin IIA heavy chain (MYH9), and through the inhibition of myosin II ATPase activity by light-insensitive S-nitro-blebbistatin. Inhibition of ROCK blocks myosin II regulatory light chain (RLC) phosphorylation, which interferes with myosin II filament assembly ^47-50^. As a result, HeLa-JWcells treated with 30 μM of Y27632 essentially lose their myosin II filaments in less than half an hour. siRNA knockdown of MYH9 also resulted in a loss of most of the myosin II filaments (fig. S3). Inhibition of myosin II ATPase activity by S-nitro-blebbistatin did not disrupt myosin II filaments ^50^, although this treatment did result in profound changes to the organization of the actomyosin cytoskeleton, including a loss of stress fibers. While treatment with either S-nitro-blebbistatin or Y27632 did not change the average filopodia length, myosin IIA knockdown resulted in moderate but statistically significant shortening of filopodia, (fig. S2), which is consistent with the results on filopodia length in wild type and myosin IIA expressing Cos-7 cells mentioned above (fig. S2). Myosin X-positive comet tails persisted at the tips of filopodia in both HeLa-JW and Cos-7cells irrespectively to inhibition or lack of myosin IIA.

Despite the morphological integrity of filopodia being preserved in myosin II inhibited or depleted cells, adhesion of filopodia to the ECM was significantly impaired. While in control HeLa-JW cells application of pulling force via fibronectin-coated bead induced sustained growth of attached filopodia accompanied by the development of up to ∼ 5 pN force, in the cells with impaired myosin II activity the filopodia detached earlier, after developing rather small forces (Fig. 3B-C, E, movies S9 B-C). This suggests filopodia are unable to establish a proper adhesion contact in the absence of active myosin IIA. We also examined the immediate effect of Y27632 during the force-induced sustained growth of filopodia. Shortly after the drug was added to the experimental chamber, filopodia indeed detached from the bead (fig. S4, movie S10).

### Interaction between actin filaments and formins is required for filopodia adhesion and myosin X localization

In myosin X-induced filopodia, the formin mDia2 is localized to the filopodia tips, and overlaps with myosin X patches (fig. S1B, upper panels). Small molecular inhibitor of formin homology domain 2 (SMIFH2) ^51^ was used to investigate the role of formins in attachment of filopodia to fibronectin-coated beads. We found that in SMIFH2 (40μM, 1hour) treated cells, adhesion of filopodia to the beads was impaired in a similar way to the adhesion of filopodia in myosin II inhibited/depleted cells. The duration of contact between the filopodia and bead was significantly shorter, and the maximal force exerted by filopodia to the bead was significantly weaker than in control cells (Fig. 3D-E, movie S9D).

While the number of filopodia in cells treated with SMIFH2 remained the same as in control cells and their mean length decreased only slightly (Fig. 4A), only about 25% of these filopodia preserved myosin X comet tails at their tips 1-2 hours following SMIFH2 addition (fig. S5; filopodia in 14 cells were scored in 2 independent experiments) despite originally being induced by over-expression of myosin X. We found that SMIFH2 induced rapid disintegration of the comet tails into myosin X patches, which rapidly moved centripetally towards the cell body (Fig. 4B, movie S12). Although such movement was occasionally observed in control cells (above), it was much more prominent in cells treated with SMIFH2, and led to the gradual disappearance of myosin X from the filopodia tips. Of note, the movement of myosin X patches in SMIFH2 treated cells occurred together with the movement of its partner VASP ^52^, another protein associated with barbed ends of actin filaments (fig. S6, movie S6).

**Fig. 4.**
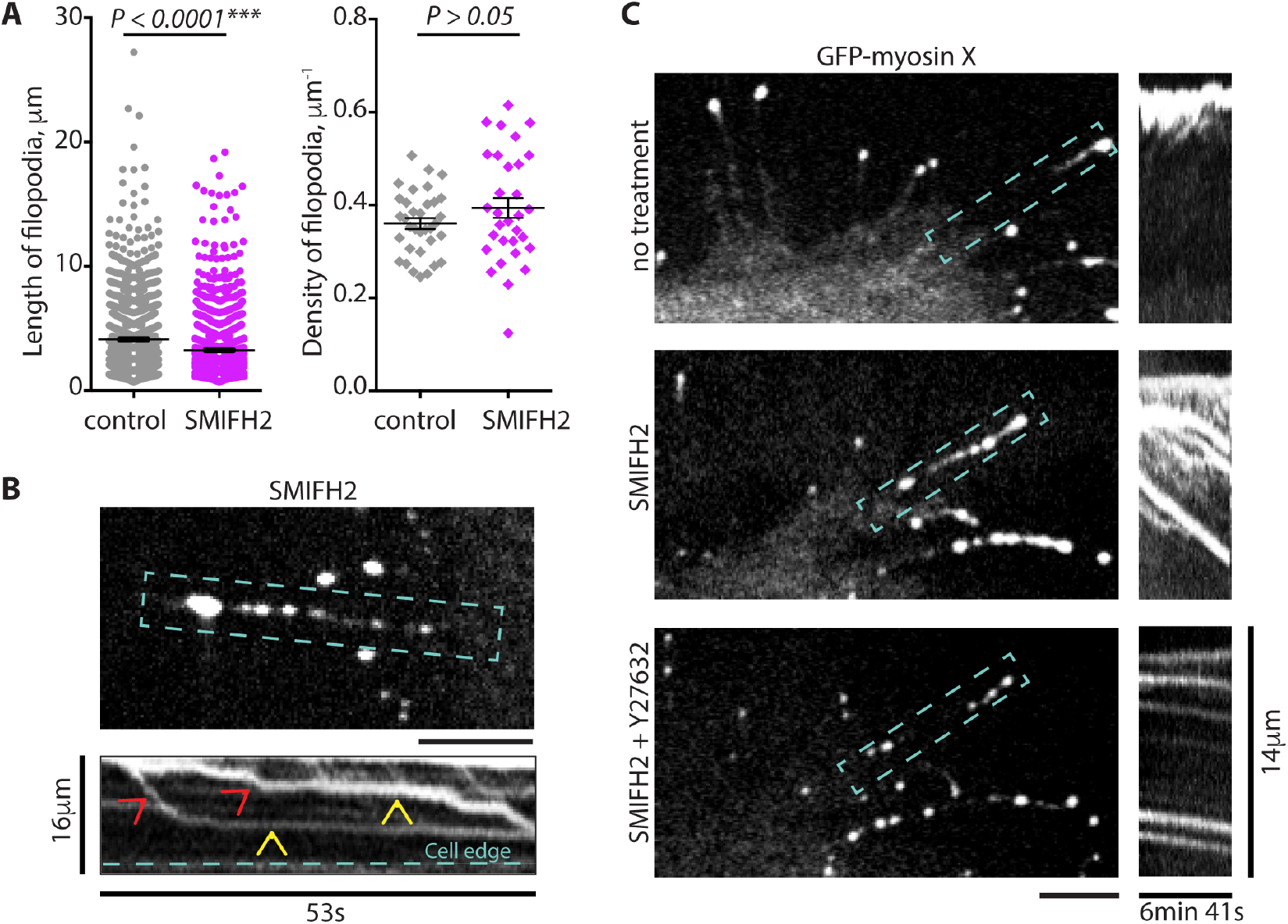
Effect of formin inhibition on filopodia growth and centripetal movement of myosin X patches. **(A)** The average length of unconstrained filopodia (left) in control HeLa-JW cells expressing GFP-myosin X (mean±SEM) was 4.1±0.1μm (n = 1710 in 34 cells), which exceeded that of SMIFH2 treated cells 3.2±0.1μm (n = 1645 in 31 cells), while the numbers of filopodia (right) per micron of cell boundary did not differ significantly (mean±SEM): 0.36±0.01 (n = 34 cells) and 0.39±0.02 (n = 31 cells), respectively. The mean values are indicated by horizontal black lines; the error bars correspond to SEMs. **(B)** Upper panel: Disintegration of the myosin X comet tail following a 2 hours exposure to 20μM SMIFH2. Numerous myosin X patches are seen in the filopodia shaft. Lower panel: A kymograph showing fast centripetal movement of the patches boxed in the upper panel towards the cell body (red arrowheads, see also movie S12). Intervals of slow centripetal movements are indicated by yellow arrowheads. **(C)** Addition of Y27632 treatment stops the movement of myosin X patches in SMIFH2 treated cells. The same filopodium is shown before SMIFH2 treatment (upper panel), 15 min after the addition of 20μM SMIFH2 (middle panel) and 15 min after subsequent addition of 30μM Y27632 (lower panel). Myosin X patches are shown in the left images (see also movies S13A-C), and kymographs representing the movement of the patches in the boxed area - in the images on the right. All images and data for analysis were obtained with SDCM. Scale bars, 5μm.

The velocity of retrograde movement of myosin X patches in filopodia of cells treated with SMIFH2 was 84 ± 22 nm/s (mean ± SEM, n = 45) vs 31 ± 5 nm/s in control cells (see the first section of Results). Such movement might be a result of the detachment of myosin X-bearing actin filaments from the filopodia tips. Once free, their subsequent retrograde movement is driven by myosin II located at the bases of the filopodia. Indeed, incubation of SMIFH2 treated cells with Y27632 efficiently stopped the retrograde movement of the myosin X positive patches, reducing their average velocity to 0.66±0.14 nm/s (mean ± SEM, n = 17), see Fig. 4C and movies S13A-C.

We also studied the immediate effect of SMIFH2 during the force-induced sustained growth of filopodia. After the drug was added, myosin X retrograde movement, cessation of the polymerization of the actin core, and the drop of the pulling force generated by a filopodium were observed (movie S11). The fibronectin-coated bead, however, remained associated with the filopodium tip via the membrane tether (movie S11). This suggests that addition of SMIFH2 induced disruption of the link between actin core and the integrin adhesion receptors, which still remain associated with the membrane tether at the filopodium tip.

To prove that SMIFH2 treatment can detach actin filaments from formin located at the filopodia tips, we performed in vitro experiments where the actin filaments were growing from immobilized formin mDia1 construct (FH1FH2DAD) in the absence or presence of SMIFH2. Following treatment with 100μM SMIFH2, a rapid decrease in the fraction of filaments remaining associated with immobilized formins under conditions of mild shear flow was observed (fig. S7). Thus, SMIFH2 treatment disrupted physical contacts between formin molecules and actin filaments. Therefore, SMIFH2-induced rapid centripetal movement of myosin X is driven by myosin II mediated pulling of actin filaments detached from the filopodia tips.

### Effect of myosin II and formin inhibition on the growth of unconstrained filopodia

In addition to the studies of filopodia growing in response to pulling forces, we examined the effects of myosin II and formin inhibition on the dynamics of free, unconstrained filopodia (Fig. 5 inset). We found that knockdown of myosin IIA and cell treatment with Y27632 or S-nitro-blebbistatin efficiently blocked growth and retraction of unconstrained filopodia, resulting in suppression of filopodia dynamics. In untreated myosin X-expressing cells, the fraction of filopodia in the “pause” state (with the growth rate between −15 and +15nm/s) was 13% (n = 194). At the same time, fractions of the “pausing” filopodia were 90% (n = 41), 80% (n = 83) and 55% (n = 42) for myosin IIA knockdown, S-nitro-blebbistatin-treated and Y27632-treated cells, respectively (Fig. 5). Similarly, the fraction of “pausing” filopodia in cells treated with the formin inhibitor SMIFH2 was 75% (n = 44) (Fig. 5)

**Fig. 5.**
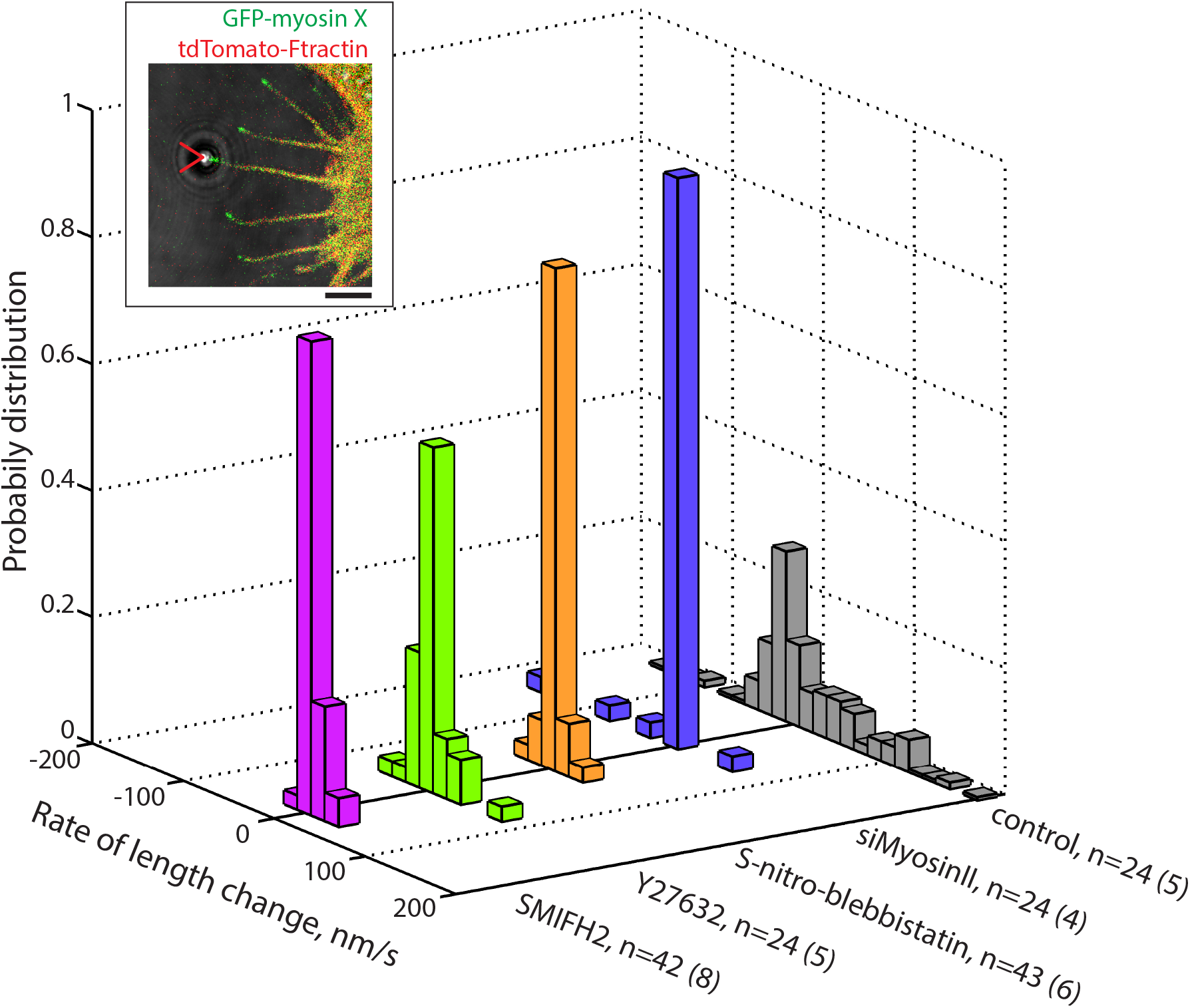
Inhibition of myosin II or formins interfere with growth of unconstrained filopodia. A graph showing the distribution of growth/retraction velocities of unconstrained HeLa-JW cells filopodia for control, myosin II siRNA knockdown, S-nitro-blebbistatin, Y27632 and SMIFH2 treatment, observed in the same experiments as those assessing the fibronectin-coated bead attachment to filopodia. *n* represents the number of processed filopodia (with number of cells in parenthesis). (Inset) A cell expressing GFP-myosin X and tdTomato-Ftractin, which is representative of those used in experiments assessing filopodia growth. The filopodium attached to the laser trapped fibronectin-coated bead is indicated by the red arrowhead. Such filopodia were excluded from the score. Scale bar, 5μm.

In this study, we have shown that filopodia adhesion to the ECM is a force dependent process. This conclusion is based on experiments in which sustained growth of filopodia was maintained by the application of pulling force at the interface between a fibronectin-coated bead, and the tip of a filopodia. With this setup, inhibition of myosin II filament formation or myosin II ATPase activity resulted in suppression of filopodia adhesion to fibronectin-coated bead. In should be noted that our experiments were performed on filopodia induced by over-expressing myosin X and, therefore, our conclusions are, strictly speaking, only valid for this class of filopodia. However, myosin X has been shown to be a universal component of filopodia ^26^, so employment of such an experimental system does not restrict the generality of our finding.

Since myosin II is located at the bases of filopodia (Fig. 2), a question requiring further clarification is how the pulling force is transmitted to the filopodia tips. Our data are consistent with the idea that this force is transmitted by the filaments of the actin core attached to formin molecules and the filopodia tips. We have shown that formin inhibition by SMIFH2 suppresses filopodia adhesion to the beads in the same manner as inhibition of myosin II. Moreover, we demonstrated that SMIFH2 treatment led to a rapid, myosin II-dependent, movement of actin filament associated proteins, myosin X and VASP, from the filopodia tips towards the cell body. Since centripetal movements of myosin X patches proceed together with the movements of photoactivated actin spots, we interpret the acceleration of myosin X/VASP movement upon SMIFH2 treatment as an evidence of actin filament detachment from formins at the filopodia tips. Indeed, *in vitro* experiments demonstrated that addition of SMIFH2 to actin filaments growing from the immobilized formins under a condition of moderate flow results in the detachment of actin filaments from the formin molecules. Together, these experiments suggest that myosin II inhibition, or inhibition of the formin-mediated association between actin filaments and the filopodia tips, make filopodia unable to form stable adhesions with fibronectin-coated beads. This in turn prevents them from growing upon force application.

To check whether adhesion and growth of filopodia require the pulling force, we compared behavior of unconstrained filopodia on rigid substrate with that on fluid supported lipid bilayer (SLB) where the traction forces cannot develop ^53^. To this end, we created a composite substrate on which rigid surface was covered by orderly patterned small islands (D = 3μm) of SLB. Both rigid and fluid areas were coated with integrin ligand, RGD peptide, with the same density (fig. S8). We found that dynamics of filopodia extended over rigid regions of this substrate was similar to that of filopodia growing on rigid fibronectin-coated substrate used in rest of experiments. At the same time, filopodia that encountered the SLB islands could not attach properly and as a result spent over such substrate significantly shorter time than over rigid area of the same geometry (fig. S9A, B, movie S14). Accordingly, the average density of filopodia tips remaining inside the SLB islands during period of observation (> 10min) was lower than that on the rigid substrate (fig. S9C). Thus, not only inhibition of myosin II or formin, but also micro-environmental conditions under which filopodia tips do not develop traction force, prevent proper adhesion of filopodia.

## DISCUSSION

In the present study, we have demonstrated that adhesion of myosin X-induced filopodia to extracellular matrix depends on the forces generated by associated myosin IIA filaments. Our interest to filopodia induced by myosin X is in part justified by the fact that this protein is overexpressed in many types of cancer cells and adhesion of myosin X containing filopodia to the matrix may play an important role in the cancer cells invasion and metastasis. ^54^

The structured illumination microscopy (SIM) revealed the presence of individual myosin IIA filaments at the base of filopodia. The force generated by one bipolar myosin IIA filament (consisting of about 30 individual myosin molecules ^55^, 15 at each side) can be estimated based on the stall force for individual myosin IIA molecule (3.4pN according ^56^) and duty ratio (5-11% according ^57^) as 2.6-5.6pN. This value is consistent with pulling forces generated by individual filopodia as measured in our experiments. In the presence of formin inhibitor SMIFH2 detaching the filaments of the actin core from formin molecules at the filopodia tips, the filaments retract centripetally towards the cell body as can be inferred from the analysis of movements of myosin X and VASP clusters associated with these filaments. Inhibition of myosin IIA filament formation by Y-27632 suppressed such retrograde movement inside the filopodia. The forces generated by myosin IIA filaments appear to be sufficient to overcome the actin filament crosslinking inside the core and generate retrograde filament movement. Altogether, these data suggest that myosin IIA-generated forces are transmitted to filopodia tips via actin cores associated with formin molecules at the tips.

Dependence of filopodia adhesion on myosin IIA-generated forces was revealed in two types of experimental systems. First, we studied the effects of application of pulling force on the filopodia. In control HeLa-JW cells, filopodia attached to optically trapped fibronectin-coated beads were growing upon slow movement of the microscope stage and developed periodic 2-5pN forces. Myosin IIA-depletion or cell treatment with inhibitors of myosin II assembly (Y-27632) or motor activity (blebbistatin) led to significant decrease of the forces exerted by filopodia and subsequent detachment of filopodia from the trapped beads. Thus, myosin IIA-generated forces are required for the adhesion of filopodia to the beads and for force-induced filopodia growth.

Second, the function of myosin IIA in the regulation of filopodia lifetime in Cos-7 cells was studied. In Cos-7 cells lacking myosin IIA, myosin X overexpression induces only unstable short living filopodia. Their lifetime, however, can be significantly increased upon expression of exogenous myosin IIA. Such increase of the lifetime can be explained by stabilization of adhesion of these filopodia to the substrate. In contrast, myosin IIB overexpression does not increase the filopodia lifetime and does not localize to filopodia bases. Importantly, even though myosin IIA mutant with compromised motor activity (myosin IIA N93K) still localizes to filopodia bases, it only slightly enhances the filopodia lifetime. Thus, the motor function of myosin IIA rather than its actin crosslinking function ^45^ is important for stabilization of filopodia adhesion.

Force-driven growth of filopodia attached to the bead is an integrin-dependent process and was not observed in experiments with integrin-independent adhesion of filopodia to beads coated by concanavalin A. At the same time, association of integrin with myosin X FERM domain was dispensable for the myosin IIA-driven augmentation of the filopodia lifetime. This suggests that potential link between integrin and actin filaments via myosin X is not critically important for filopodia adhesion stabilization.

A major linker between integrin and actin filaments, talin, has been detected at the filopodia tips in this and other publications ^30^. Previously, it was established that force-driven unfolding of talin facilitates interaction of talin with another adhesion complex component, vinculin, resulting in reinforcement of the association between talin and actin filaments ^58–61^. The question whether this mechanism is applicable to filopodia adhesion reinforcement requires additional studies. While vinculin enrichment was detected in shafts but not tips of filopodia ^62^, a RIAM protein that competes with vinculin for talin-binding and localizes to filopodia tips ^30, 63^ may be involved in force dependent filopodia adhesion. RIAM binds Ena/VASP and profilin^64^ and could recruit these actin polymerization-promoting proteins to the filopodia tips.

Another mechanism of myosin II-dependence of filopodia adhesion and growth might involve formin-driven actin polymerization known to be a major factor in filopodia extension ^15–17, 19–21^. Recent studies demonstrate that formin-driven actin polymerization can be enhanced by pulling forces ^65–68^. Thus, myosin II-generated force transmitted via actin core to formins at the filopodium tip can stimulate actin polymerization, promoting filopodia growth. Polymerization of actin could also be important for recruitment of new adhesion components to the filopodia tips and adhesion reinforcement ^69^.

Filopodia adhesion is tightly associated with filopodia growth and shrinking. In our experiments, inhibition of myosin II and formins not only suppressed filopodia adhesion but also resulted in reduction of motility of filopodia along the substrate. During pulling-induced growth of bead-attached filopodia, periods of filopodia elongation alternate with periods of growth cessation accompanied by increase of the pulling force. Notably the periods of force development preceded the periods of filopodia elongation. Thus, force developed during growth cessation may trigger the subsequent filopodia growth. Similarly, the growth of unconstrained filopodia along rigid substrate can proceed via periods of attachment, development of force, and consequent filopodia elongation ^41^. Inhibition of force generation or transmission suppresses such dynamics. Interestingly, in filopodia of other types the attachment of filopodia tips to the substrate instead of further elongation led to formation of associated lamellipodia ^70^. It is not yet known, whether this switch is also mediated by myosin II-generated force.

Our finding that filopodia adhesion and growth are both is force-dependent explains how filopodia could respond differently to substrates of varying stiffness. On a stiff substrate, the force generated by myosin II and applied to the adhesion complex will develop faster than on a compliant substrate ^71^. Accordingly, filopodia adhesion should be more efficient on stiff substrates than on compliant substrates. Indeed, we showed that the contacts of filopodia with RGD ligands associated with fluid membrane bilayer were less stable than with the areas of rigid substrate covered with RGD of the same density. These considerations can explain involvement of filopodia in the phenomenon of durotaxis ^39^, a preferential cell movement towards stiffer substrates ^72^. This may provide a mechanism to rectify directional cell migration.

Orientation based on filopodia adhesion is characteristic for several cell types, in particular for nerve cells. The growth cones of most neurites produce numerous filopodia, and the adhesion of these filopodia can determine the direction of neurite growth ^73, 74^ Interestingly, the filopodia-mediated traction force in growth cones is myosin II-dependent ^75^ and application of external force can regulate the direction of growth cone advance ^76^. The results from these experiments can now be explained by preferential adhesion/growth of filopodia, which experience larger force. The mechanosensitivity of filopodia adhesion provides a mechanism of cell orientation that complements that mediated by focal adhesions. Focal adhesions are formed by cells attached to rigid two-dimensional substrates, whereas filopodia adhesion can be formed by cells embedded in three-dimensional fibrillar ECM network. Thus, further investigation of filopodia mechanosensitivity could shed a new light on a variety of processes related to tissue morphogenesis.

## METHODS

### Cell culture and transfection

Hela-JW, a subline of a HeLa cervical carcinoma cell line derived in the laboratory of J. Willams (Carnegie-Mellon University, USA) on the basis of better attachment to plastic dishes ^77^, was obtained from the laboratory of B. Geiger ^78^. Cos-7 (African green monkey fibroblast-like cell line, was obtain from ATCC (ATCC^®^ CRL-1651™). The cells from both cell lines were grown in DMEM-Dulbecco’s Modified Eagle Medium supplemented with 10% fetal bovine serum (FBS), 100 U ml^−1^ penicillin/streptomycin, 2 mM glutamine and 1 mM sodium pyruvate at 5% CO_2_ at 37 °C. HeLa-JW cells were transfected with DNA plasmids by electroporation (two pulses of 1005 V for 35 ms) using Neon transfection system (Thermofisher), while Cos-7 cells – by jetPRIME transfection reagent (Polyplus) according to manufacturer protocol. Expression vectors encoding the following fluorescent fusion proteins were used: GFP-myosin X and GFP-Myo10 ΔFERM ^40, 44^ (gift from R. Cheney, University of North Carolina Medical School, USA), reduced size myosin X construct (without 3’UTR) derived from GFP-myosin X was sub-cloned into mApple-C1 cloning vector backbone (M. Davidson collection, Florida State University, USA, kindly provided by P. Kanchanawong, MBI, NUS, Singapore), tdTomato–Ftractin ^79^ (gift from M. J. Schell, Uniformed Services University, Bethesda, Maryland, USA), mCherry-Utrophin ^80, 81^ was kindly provided by W. Bement (University of Wisconsin, Madison, USA), mTagBFP-Lifeact was obtained from M. Davidson collection, Florida State University, USA (Addgene), myosin regulatory light chain (RLC)-GFP^82^ (gift from W. Wolf and R. Chisholm, Northwestern University, Chicago, Illinois, USA), GFP-myosin IIA heavy chain was sub-cloned from pTRE-GFP-NMHCIIA (from R. Adelstein, NHLBI, NIH, USA) into pEGFP-C3 by MluI & HindIIIcloning sites (cloned by M. Tamada, M. Sheetz laboratory), GFP-myosin IIA N93K ^49^ was kindly provided by Dr. Vicente-Manzanares (Universidad Autónoma de Madrid, Madrin, Spain), GFP-myosin IIB, mCherry-VASP and mCherry-talin (M. Davidson collection, Florida State University, kindly provided by P. Kanchanawong, MBI, NUS, Singapore), full-length mDia2 was sub-cloned from GFP-C1-mDia2 ^83^ (gift from S. Narumiya, Faculty of Medicine, Kyoto University, Japan) into mCherry-C1 vector(Clontech), PAmCherry-b-actin was kindly provided by V. Verkhusha (Albert Einstein College of Medicine, NY, USA). All cell culture and transfection materials were obtained from Life Technologies.

### Live cell imaging and confocal microscopy

Following electroporation, cells were seeded at a density of 2 × 10^4^ cells ml^−1^ in 2ml onto 35mm glass based dishes with 12 or 27 mm bottom base cover glass #1 in diameter (Iwaki, Japan) coated with 10 μg ml^−1^ fibronectin (Calbiochem) for 20 min. Cells were imaged in Leibovitz’s L-15 medium without Phenol Red containing 10% FBS at 5% CO_2_ at 37 °C. Snapshot or time-lapse images were acquired with a spinning-disc confocal system (PerkinElmer Ultraview VoX) based on an Olympus IX81 inverted microscope, equipped with a 100 × oil immersion objective (1.40 NA, UPlanSApo), an EMCCD camera (C9100-13, Hamamatsu Photonics), and Volocity control software (PerkinElmer).

We perform photoactivation experiments by activation of a defined region inside filopodia using blue diode laser (405 nm, 100mW). Photoactivation and livecell imaging were performed on CSU-W1 spinning disk confocal system on Nikon Eclipse Ti-E inverted microscope with Perfect Focus System, controlled by MetaMorph software (Molecular device) supplemented with a 100x oil 1.45 NA CFI Plan Apo Lambda oil immersion objective and sCMOS camera (Prime 95B, Photometrics).

For structured illumination microscopy (SIM), two types of equipment were used: (1) Live SR (Roper Scientific) module on Nikon Eclipse Ti-E inverted microscope (specifications of setup described above), (2) Nikon N-SIM microscope, based on a Nikon Ti-E inverted microscope with Perfect Focus System controlled by Nikon NIS-Elements AR software supplemented with a 100x oil immersion objective (1.40 NA, CFI Plan-ApochromatVC) and EMCCD camera (Andor Ixon DU-897). For life cell imaging the samples were mounted in a humidified cell culture chamber and maintained at 37 °C with 5% CO_2_. All SIM images with obtained with system (1) except images on Fig. 2A and B (upper panel), which were obtained with set up (2).

### Transfection of siRNA and immunoblotting

Cells were seeded into a 35 mm dish on day 0 and transfected with 100μM of MYH9 ON-TARGET plus SMART pool siRNA (L-007668-00-0005, Dharmacon) using Screenfect^TM^A (WAKO, Japan) on day 1. Control cells were transfected with scrambled control ON-TARGET plus Non-targeting pool siRNA (D-001810-10, Dharmacon). Transfection of plasmid GFP-myosin X and tdTomato-Ftractin was performed in the evening of the day 1 using Jet Prime transfection reagent (Polyplus) and cells were imaged on day 2. For assessment of myosin IIA heavy chain expression, transfected cells were lysed in RIPA buffer on day 2 (exactly 24 hours following siRNA transfection) and analyzed by Western blotting with primary rabbit antibodies to the myosin IIA tail domain (M8064, Sigma-Aldrich, dilution 1:1000); staining of α-tubulin with monoclonal DM1A antibody (T6199, Sigma-Aldrich, dilution 1:5000) was used as a loading control. HRP-conjugated anti-rabbit IgG (Bio-Rad, 1706515, dilution 5000) and anti-mouse IgG (A4416, Sigma-Aldrich, dilution 1:10000) were used as secondary antibodies, respectively.

### Immunofluorescence antibody staining

Anti-myosin IIA tail domain (M8064, Sigma-Aldrich, dilution 1:800), and 405 Alexa-Fluor-conjugated secondary antibodies (A31556, Molecular Probes, dilution 1:200). Cells were pre-fixed by addition of warm 20% PFA (Tousimis) into medium (to make 8% solution) and subsequent 15 min incubation at room temperature. This was followed by fixation and permeabilization by 3.7% PFA, 0.2% glutaraldehyde and 0.25% Triton X-100 in PBS for 15 min. The fixed cells were then washed two times with PBS and blocked with 5% bovine serum albumin (BSA) for 30 min. The cells were then stained with primary antibodies overnight at 4 °C, and incubated with secondary antibodies for 1h at room temperature.

### Drug treatment

For formin drug inhibition studies, cells were incubated with 20 or 40μM SMIFH2 (4401, TOCRIS, UK) in serum containing DMEM for 1-2h at 5% CO_2_, 37 °C. In *in vitro* experiments, SMIFH2 (340316-62-3, Sigma-Aldrich) was used at a concentration of 100μM (see below). For myosin II inhibition studies, 30μM or 50μM Rho-kinase (ROCK) inhibitor Y27632 (Y0503, Sigma-Aldrich) or 20μM S-nitro-blebbistatin (13013-10, Cayman Chemicals) was added for 10-20 min before the experiments or directly during observations. All inhibitors remained in the medium during the entire period of observation.

### Optical tweezers and data acquisition

All experiments involving filopodia pulling were carried out on a Nikon A1R confocal microscope adapted for the use of laser tweezers. 2.19 μm diameter polystyrene beads (PC05N, Bangs Laboratories) were coated with fibronectin (341635, Calbiochem) according to a previous protocol ^84^ For bead trapping we used an infrared laser (λ = 1064nm, power 0.5-1W, YLM-5-LP-SC Ytterbium Fiber Laser, IPG photonics).

To determine the forces, F, applied to the bead by the optical trap, we measured the displacement of the bead from the trap center, Δx, and then knowing the stiffness of the trap, k, the force was calculated as: F = kΔx. The trap stiffness was calibrated using the equipartition method ^85^ by tracking the fluctuations of a bead trapped by optical tweezers, using an Andor Neo sCMOS camera, at 100fps. The displacement of the beads from the center of the optical trap in confocal microscopy observations was monitored using piA640-210gm camera (Basler) at 0.5-1fps and Metamorph software for tracking. The smallest detectable bead displacement was ∼5 nm, corresponding to the smallest force measured of ∼0.04 pN.

For laser trap experiments, HeLa-JW cells transfected by electroporation with GFP-myosin X and tdTomato-Ftractin, were seeded at a density of 2 × 10^4^ cells ml^−1^ in 750μl onto chambered #1 borosilicate cover glasses (155383, Lab-Tek) coated with fibronectin (341635, Calbiochem) by incubation in 1ng ml^−1^ solution in PBS for 20min at 37.0 °C. A reduced concentration of fibronectin (compared to that used for regular cell observations) was used to prevent the beads sticking to the cover slip. Chambers with cells were mounted on P-545.3R7 stage equipped with E-545 Plnano piezo controller (Physik Instrumente), which was moved in order to generate the pulling force between filopodia and trapped beads. The velocity of the stage movement was maintained in a range 10-20 nm/s by PIMikroMove 2.15.0.0 software. The specimen were incubated in a custom-built microscope hood at 37.0 °C, 5% CO_2_ humidified environment. Simultaneously with pulling, the cells were imaged using lasers λ = 488nm and 561nm for excitation of GFP and tdTomato, respectively. Collected experimental data were processed by particle tracking algorithms of the Metamorph software.

### *In vitro* assay for formin processivity in the presence of SMIFH2

We used a microfluidics based assay to assess formin processivity in the course of formin-driven actin polymerization. The method has been described in more detail elsewhere ^66^. Briefly, formin construct Snap-mDia1(FH1FH2DAD)-6xHisTag was specifically anchored to the bottom surface of a microchamber, using a biotinylated pentaHis antibody (Qiagen) and streptavidin to bind to a biotin-BSA-functionalized glass surface which was further passivated with BSA. The microfluidics PDMS chamber height was 60 μm. Immobilized formin constructs were allowed to nucleate actin filaments by exposing them for 30s to a 2 μM solution of 20% Alexa488-labeled at Lys^328^ actin ^86^ and 0.4μM profilin in a buffer containing 10 mM Tris-HCl (Euromedex) pH 7.8, 1 mM MgCl2 (Merck), 200 μM ATP (Roche), 50 mM KCl (VWR Chemicals) and supplemented with 5 mM DTT (Euromedex), 1 mM DABCO (1,4-diazabicyclo[2.2.2]octane) to reduce photobleaching. Recording of actin filament elongation was started after the buffer was changed to contain 1 μM unlabeled actin and 4 μM profilin, with or without 100 μM SMIFH2 (Sigma-Aldrich). The microfluidics flow was kept low enough to ensure that the viscous drag had no impact on filament detachment from formin. Images were acquired every 10 s on a Nikon Ti-E microscope using a 60x objective and a Hamamatsu Orca Flash 4.0 V2+ sCMOS camera. Actin was purified from rabbit muscle ^87^. Recombinant Profilin I and Snap-mDia1(FH1FH2DAD)-6xHisTag were expressed in *E. Coli* and purified ^88^.

### Filopodia density and length measurement; tracking filopodia tips

Cells labeled with GFP-myosin X, tdTomato-Ftractin or mCherry-Utrophin were imaged in 488nm (green) and 594nm (red) channels, simultaneously. Cell segmentation was performed using an in-house algorithm implemented as a Fiji macro. The segmented cell was analyzed using MATLAB software CellGeo ^89^ to estimate filopodia density and their length. First, each data set, which corresponded to 10 frames per movie taken at 1-2 fps at x100, was averaged over time to generate a smooth image for each channel. The averaged images from the green and red channels were then summed to produce an enhanced image for segmentation. To optimize the segmentation results, filopodia segmentation and cell body segmentation was conducted in two separate steps. For filopodia segmentation, a background subtraction procedure with a rolling ball of 20-pixel radius was applied first, followed by Triangle auto-thresholding (Fiji Auto Threshold v1.16.1). For cell body segmentation, Gray Morphology open operator with a circle of 5-pixel radius was used (Fiji Morphology) to remove filopodia before applying Li auto-thresholding (Fiji Auto Threshold v1.16.1). The final segmentation result was obtained by combining the masks from the above two steps. After segmentation, the BisectoGraph module of CellGeo ^89^ was used to partition each cell into the cell body and individual filopodia, based on the parameters called critical radius (roughly the half of maximal filopodia width) and minimum filopodia length. In our analysis, a critical radius of 0.7μm and a minimum filopodia length of 1.5μm were used. Note that, our definition of filopodia length slightly differed from that used in original paper by ^89^. Namely, in our analysis the filopodia length is defined to be the distance from the filopodia tip to the cell body boundary. The MATLAB code for computing this length was kindly provided by D. Tsygankov (Georgia Institute of Technology, Atlanta, GA, USA). For filopodia density quantification, the perimeter of the cell body was measured and the number of filopodia per unit of length (μm^−1^) was computed.

To track the filopodia tips, the Imaris 8.3.1 spots tracking procedure was used. The filopodia tips were identified using segmentation procedure described above followed by MATLAB binary morphology function bwmorph with ‘skel’ and ‘endpoints’ operations.

Unconstrained filopodia elongation and shrinking rates were obtained by linear regression fitting of the filopodia tips trajectories at the regions corresponding to the periods of filopodia persistent growth or shrinking, respectively.

### Supported lipid bilayer micro patterns chamber and filopodia tips trajectory analysis on rigid and fluid substrates

The cleaned glass coverslips were coated with PLL-g-PEG-biotin (Susos) for two hours followed by UV etching using 3 μm circular shape arrays of chromium (Cr) photomask ^90^, which resulted in removal of PLL-g-PEG-biotin polymers on the etched circular regions. After multiple rinses with UHQ water, phospholipid vesicles were introduced to the surface for 5 min, allowing the self-assembly of lipid bilayer on the etched glass surfaces. Phospholipid vesicles were synthesized using the existing protocol ^90^. Specifically, 98 % of DOPC (1,2-dioleoyl-sn-glycero-3-phosphocholine) was mixed with 2 % of biotin-DOGC (1,2-dioleoylsn-glycero-3-[(N-(5-amino-1-carboxypentyl)iminodiacetic acid)-succinyl]) biotin lipids. The lipid solution was further diluted with Tris buffered saline (TBS, Sigma-Aldrich) at a ratio of 1:1 before introduction. The glass coverslips were assembled into the donut shape chamber at the UHQ water reservoir. The hybrid substrate was then incubated with 0.1 % bovine serum albumin (BSA, Sigma Aldrich) for two hours to reduce nonspecific protein absorption. Next, 1 μg/ml of Dylight-405 NeutrAvidin (Thermo Fisher) was introduced to the chamber for one hour, and then incubated with 1ug/ml of RGD-biotin (Peptides International) for one hour. Multiple rinses with UHQ were applied after each step. A fluidity test of RGD molecules at the surface of supported lipid bilayer (SLB) was performed by fluorescence recovery after photobleaching. The concentration of RGD on both SLB and PLL-g-PEG polymer was similar, based on the estimated fluorescence of Dylight-405 NeutrAvidin (fig. S8).

HeLa-JW cells expressing a myosinX-GFP chimera were introduced to the chamber, allowing cells to adhere to and spread on the RGD-coated hybrid substrate. Cells were visualized using a charge coupled EMCCD camera (Photometrics) coupled with total internal reflection microscopy (TIRF) with 100x objective (1.5 NA, Nikon). The chamber was maintained at 37°C during observation. The time-lapse video was recorded at varying intervals over a range of 3 – 5 s.

The GFP-Myosin X cluster at the filopodia tip was tracked by using a cross-correlation single particle tracking method ^91^. Obtained trajectories were analyzed using the “inpolygon” matlab function to find trajectory segments that crossed SLB circular islands and computer drawn islands in the center of the SLB pattern. The time intervals between the initial and end points of the segments were then used to estimate the average filopodia tip dwell time on the SLB and computer drawn circular islands. In addition, the ratio between the number of filopodia tip trajectories remaining inside rigid and fluid circles relatively to the total number of trajectories in the circles during the period of observation was calculated to characterize the filopodia adhesion preferences to rigid and fluid substrates (Fig. 6). In this analysis, we used only trajectories that spanned more than 5 frames.

### Statistics and reproducibility

Prism (GraphPad 6.0 Software) was used for statistical analysis. Each exact n value is indicated in the corresponding figure or figure legend. The significance of the differences (P value) was calculated using the two-tailed unpaired Student’s *t*-test.

## Acknowledgments

Encouraging and stimulating discussions with Drs. D. Bray (University of Cambrige, UK), M.M. Kozlov (Tel Aviv University, Israel) are much appreciated. We are grateful to Dr. T. Kachanawong (MBI, Singapore) for providing genetic constructs, Dr. D. Kovar (University of Chicago, IL, USA) for a sample of SMIFH2 inhibitor, and Dr. Tsygankov (Georgia Tech, USA) for providing code for filopodia length computation. We thank the Protein Cloning and Expression Core facility of the MBI for help with sub-cloning of mCherry-mDia2 and mApple-myosin X. We also thank Dr. F. Margadant and Lau Wai Han (MBI Microscopy Core facility) and Dr. V. Vyasnoff (MBI, Singapore) for their kind help with the optical tweezers setup. This research has been supported by the National Research Foundation Singapore, Ministry of Education of Singapore, Grant R714006006271 & R714019006271 (awarded to A.D.B.), Grant MOE2012T31001 (awarded to Y.J.) and BMRC Grant A*Star-JST 1514324022 (awarded to A.D.B.). As well we thank S. Wolf and A. Wang (Science Communication MBI, Singapore) for excellent editorial help.

---

**Fig. S1.**
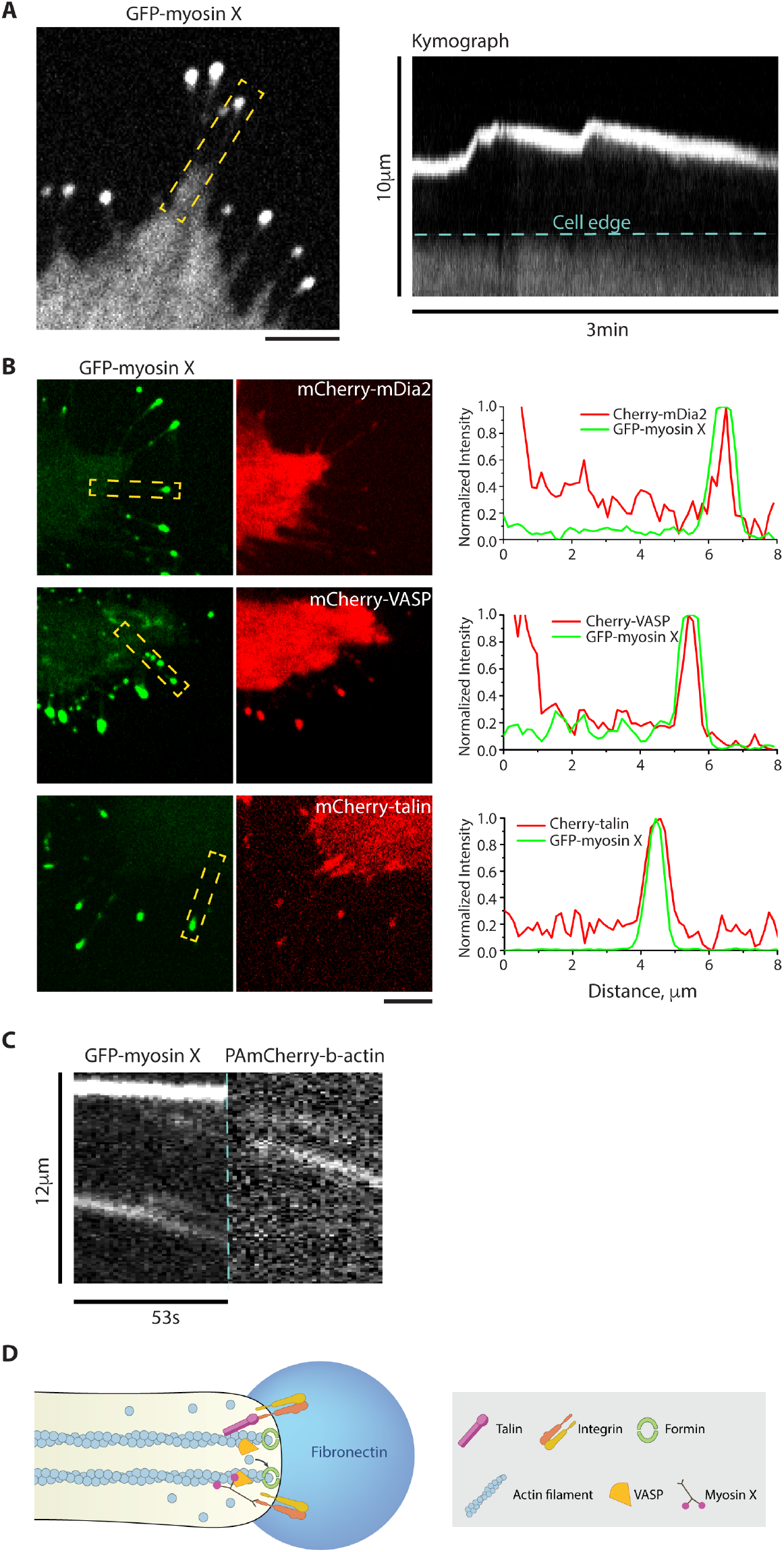
Dynamics and composition of filopodia induced by myosin X expression. **(A)** Left: Filopodia from HeLa-JW cells expressing GFP-myosin X. Myosin X positive “comet tails” are seen at the tips of filopodia. Right: A kymograph showing the growth of a GFP-myosin X labeled filopodium indicated by dashed box in the left image. Note that periods of fast growth alternate with periods of slow shrinking. See also movies S1. **(B)** Filopodia tips labeled with myosin X are enriched with mDia2, VASP and talin. Left: Images of filopodia in cells co-expressing GFP-myosin X with mCherry fusion constructs of mDia2, VASP and talin respectively. Right: Line scans of the fluorescence intensities through the filopodia indicated by dashed boxes in the left images. Intensities of the myosin X, mDia2, VASP and formin were normalized to their maximal values at the filopodia tips. Scale bars, 5μm. **(C)** Kymograph analysis of the retrograde movement of myosin X and photoactivated actin in the same filopodia. Left panel corresponds to GFP-myosin X and shows immobile tip of the filopodium (upper line) and retrograde movement of myosin X (tilted line). Right panel represents the retrograde movement of photoactivated PAmCherry-b-actin (tilted line) in the same filopodia. Note that the rate of retrograde movements of myosin X and actin are similar and were equal to 37nm/s. See also movie S7. **(D)** A cartoon depicting the protein composition at the tip of filopodium. Actin filaments are connected with integrin receptors via talin and polymerized by formin family. VASP proteins are located at the tip. Myosin X is interacting with actin filaments, integrins and VASP. Images show in panels (A) and (B) were obtained with SDCM; (C) was obtained with SIM.

**Fig. S2.**
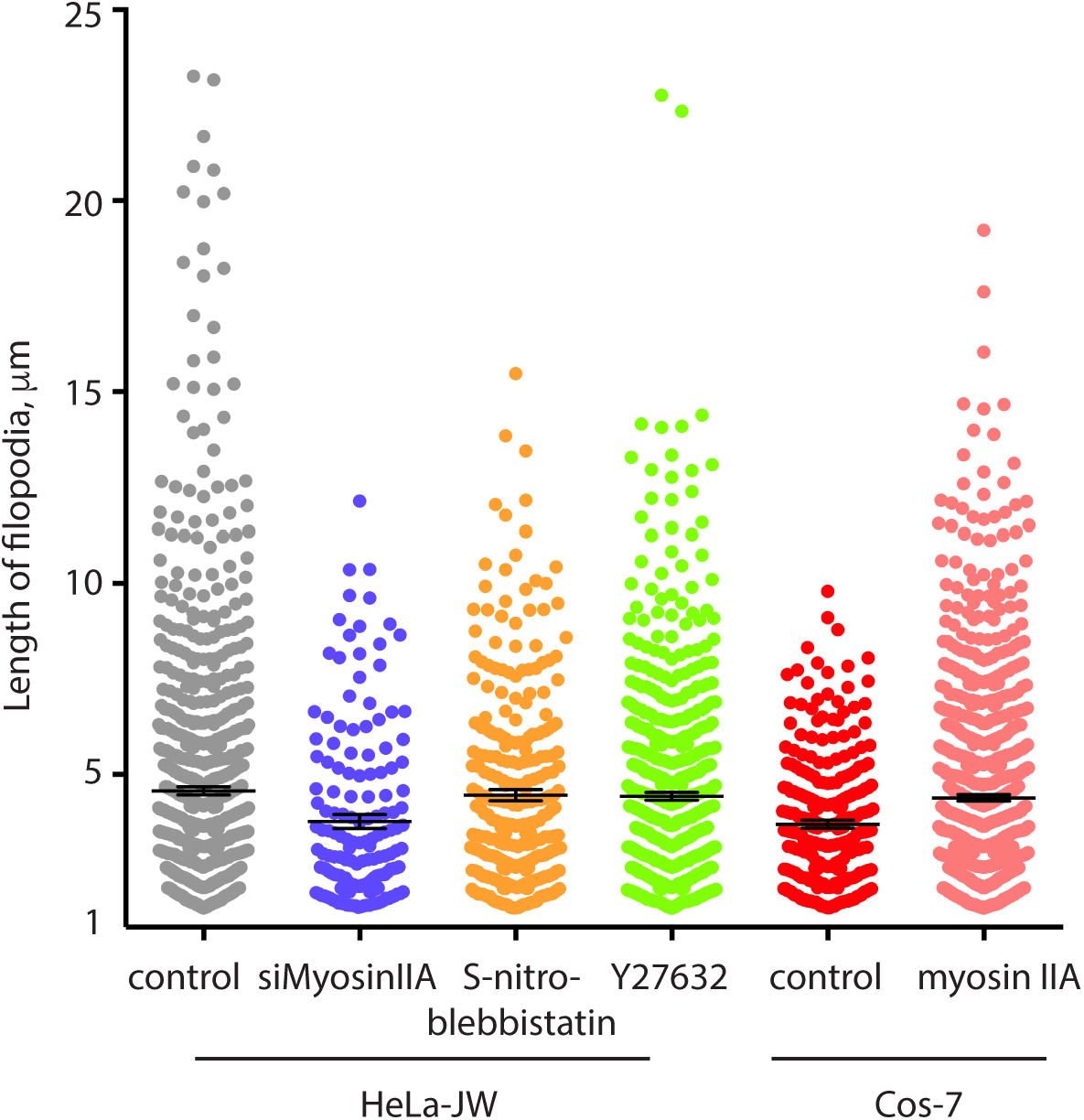
Length of myosin X-induced filopodia in HeLa-JW and Cos-7 cells. From left to right: Scattered dot plots representing the lengths of filopodia in HeLa-JW cells expressing GFP-myosin X untreated (control) and myosin IIA-depleted cells, as well as treated with S-nitro blebbistatin (20μM, 1hour) and with Y27632 (30μM, 1 hour). The last two plots represent control myosin X-expressing Cos-7 cells and Cos-7 cells co-transfected with myosin X and myosin IIA. The symbols correspond to individual filopodia. The mean values are indicated by horizontal black lines; the error bars correspond to SEMs. The mean lengths of control myosin X-induced filopodia, and filopodia from myosin IIA siRNA-, S-nitro-blebbistatin-, and Y27632-treated cells, were (mean±SEM) 4.6±0.1μm (n = 922, 38 cells), 3.8±0.2μm (n = 160, 18 cells), 4.5±0.1μm (n = 299, 24 cells), 4.4±0.1μm (n = 656, 27 cells), respectively. The mean lengths of filopodia in myosin X-transfected control Cos-7 cells and Cos-7 cells co-expressing myosin X and myosin IIA were: 3.7±0.1μm (n = 269, 11 cells) and 4.4±0.1μm (n = 950, 32 cells) respectively.

**Fig. S3:**
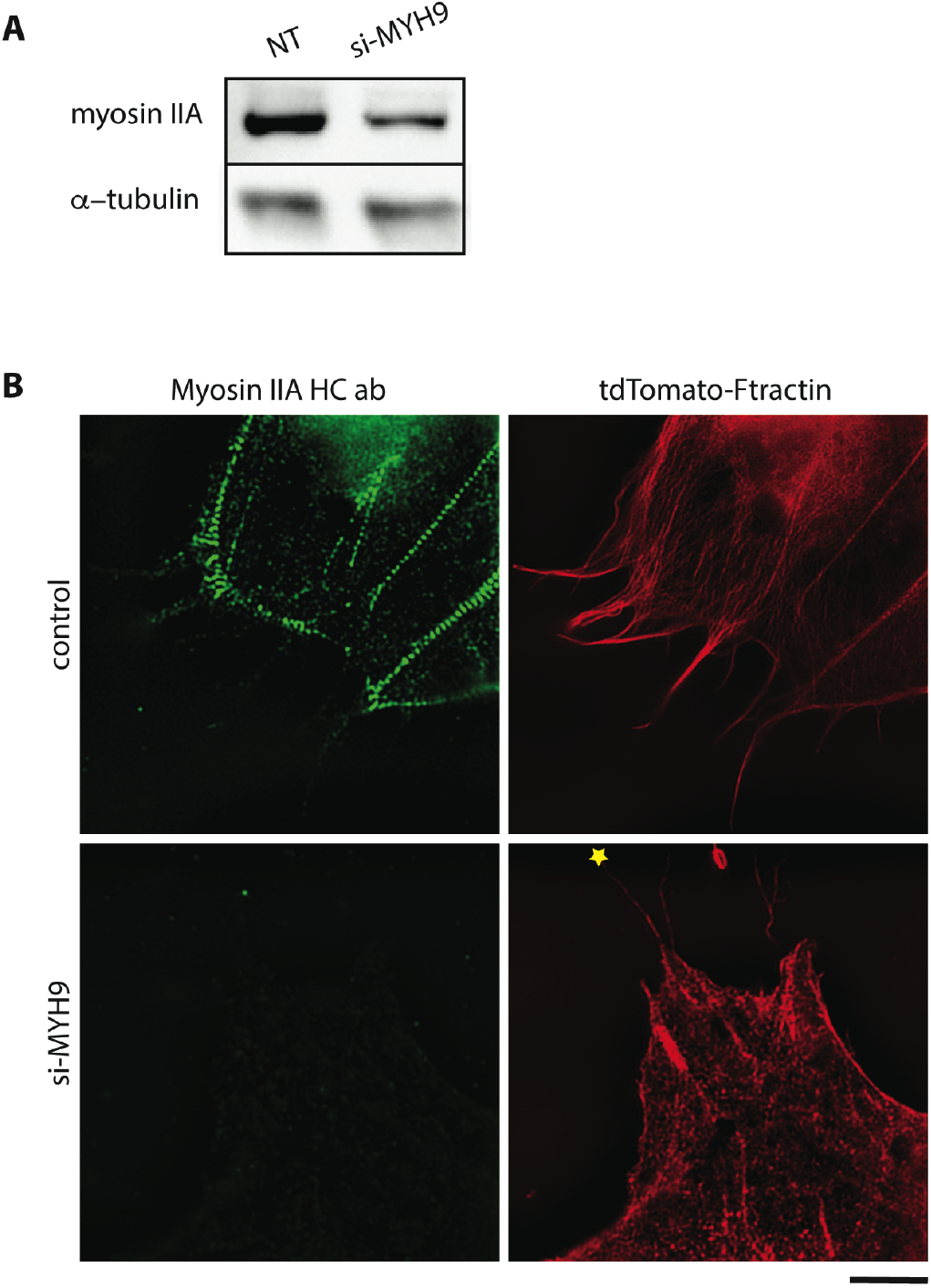
siRNA knockdown of myosinIIA. **(A)** Immunoblots of myosin IIA in non-targeted and knockdown HeLa-JW cells. α-tubulin was used as loading control. **(B)** Myosin knockdown cells did not contain either myosin IIA (left bottom) or prominent actin stress fibers (right bottom), but still contained filopodia. The filopodium attached to the bead, which was stretched in one of optical trap experiments, is indicated by asterisk. Images were obtained with SDCM. Scale bar, 10μm.

**Fig. S4.**
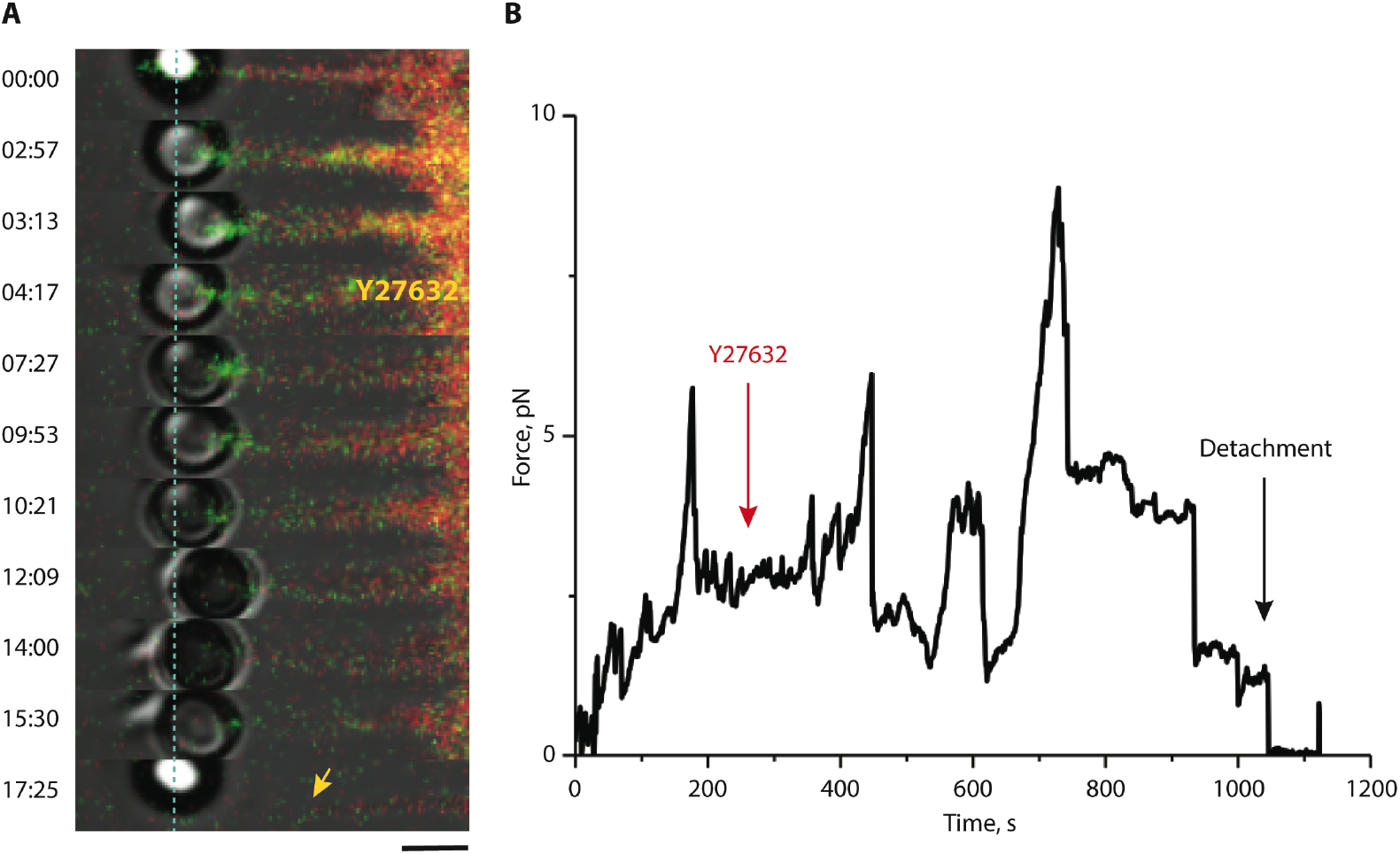
Immediate effects of ROCK inhibitor, Y27632, and formin inhibitor, SMIFH2, on force-induced filopodia growth and adhesion. **(A)** A fibronectin-coated bead, which was trapped by laser tweezers, was attached to the filopodium tip of HeLa-JW cell. Filopodium growth was then induced through the generation of pulling force, which resulted from the movement of the microscope stage, as shown in Fig. 1. At about 4 min following the start of the stage movement, 30μM of Y27632 was added, and this resulted in bead detachment at about 17 min. The position of the trapped bead during the filopodium growth is shown. The intensity of actin labeling was relatively low and apparent “disappearance” of actin in the late frames was a result of photobleacning. See also movie S10. **(B)** Deflection of the bead position from the trap center was used to calculate the forces exerted by the filopodium on the bead. Scale bar, 2μm. (C) **(D)**

**Fig. S5.**
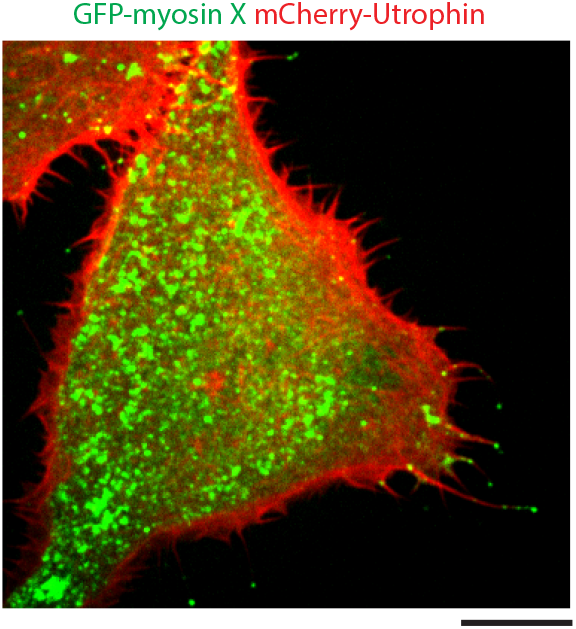
Effect of formin inhibitor SMIFH2 on myosin X induced filopodia. Image of HeLa-JW cell, labeled with GFP-myosin X and Cherry-Utrophin, that was treated with 20μM SMIFH2 formin inhibitor for 2 hours. Note that the majority of the filopodia do not contain myosin X. The image was obtained using SDCM. Scale bar, 10μm.

**Fig. S6.**
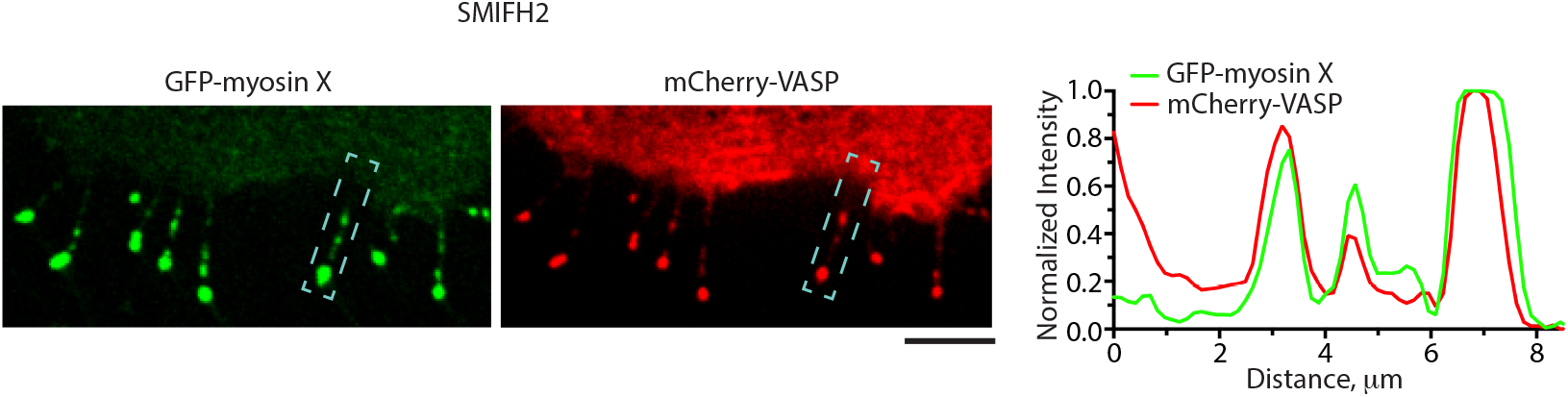
Localization of VASP at myosin X patches in cells treated with formin inhibitor. Distribution of GFP-myosin X (green, left panel) and Cherry-VASP (red, central panel) in the same HeLa-JW cell 1.5h after addition of 20μM SMIFH2. Right panel: the line scans through the boxed area, showing the co-distribution of myosin X and VASP. Fluorescent intensities are normalized to their maximal values. Scale bar, 5μm. Images were obtained with SDCM.

**Fig. S7.**
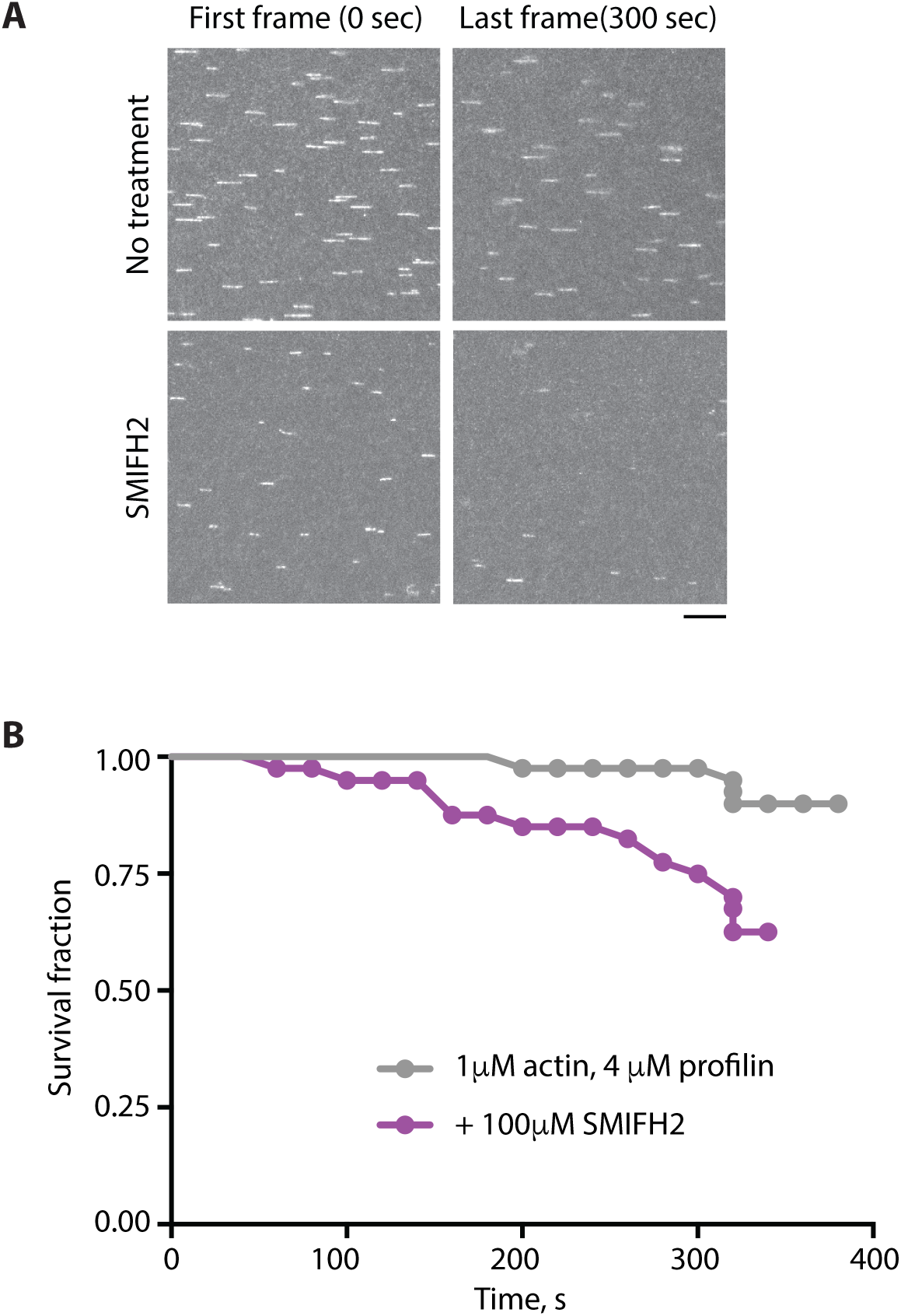
SMIFH2 enhances the detachment of formins from actin filaments in vitro. The constitutively active mDia1 formin construct (FH1FH2DAD) was anchored to the glass surface of a microfluidic chamber by one of their FH2 domains using an anti-His antibody (as in Jegou et al. 2013, see Materials and Methods). Actin filaments were grown first in the presence of Alexa488-labelled actin to form fluorescent segments at the tips of actin filaments. Once the formin had nucleated a filament, it was exposed, from time zero onward, to 1μM unlabeled actin and 4μM profilin, in absence or presence of 100 μM SMIFH2. Panel (A) shows frames corresponding to the beginning (time zero) and 300 sec following addition of unlabeled actin into the flow chamber in the presence or absence of SMIFH2 inhibitor in solution. The epifluorescence images of the labeled segments of actin filaments are seen. The number of filaments decreased with time due to their detachment from immobilized formin molecules. The time at which each filament detached from its formin was recorded and the survival fraction of the filaments at each time point was calculated. (B) SMIFH2 enhanced the filament detachment rate by an order of magnitude without affecting the formin mediated elongation rates of the filaments (35 ± 6 vs. 32 ± 8 subunits/s, n = 40 filaments without and with SMIFH2, respectively). Scale bar, 10μm.

**Fig. S8.**
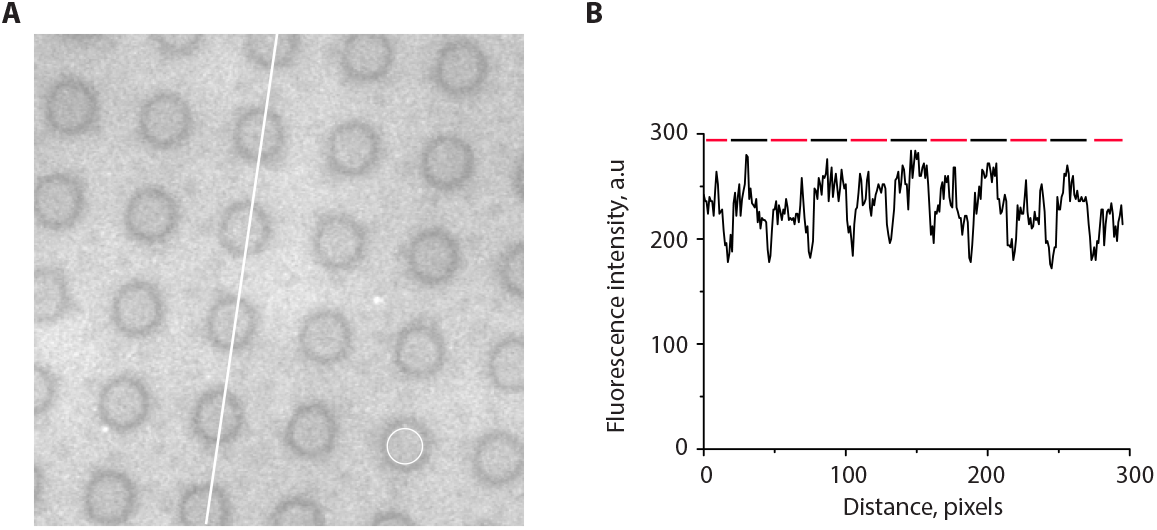
**(A)** A fluorescence image of a hybrid substrate containing a RGD ligand which is tethered to PLL-PEG polymer (background) and a supported lipid bilayer (SLB) surface (circles) via a biotin-Dylight-405 NeutrAvidin conjugation. **(B)** RGD intensity profile along the white line shown in (A). The red and black lines at the top of the curve mark the SLB and PLL-PEG regions, respectively, showing a similar concentration of RGD ligands in both regions.

**Fig. S9.**
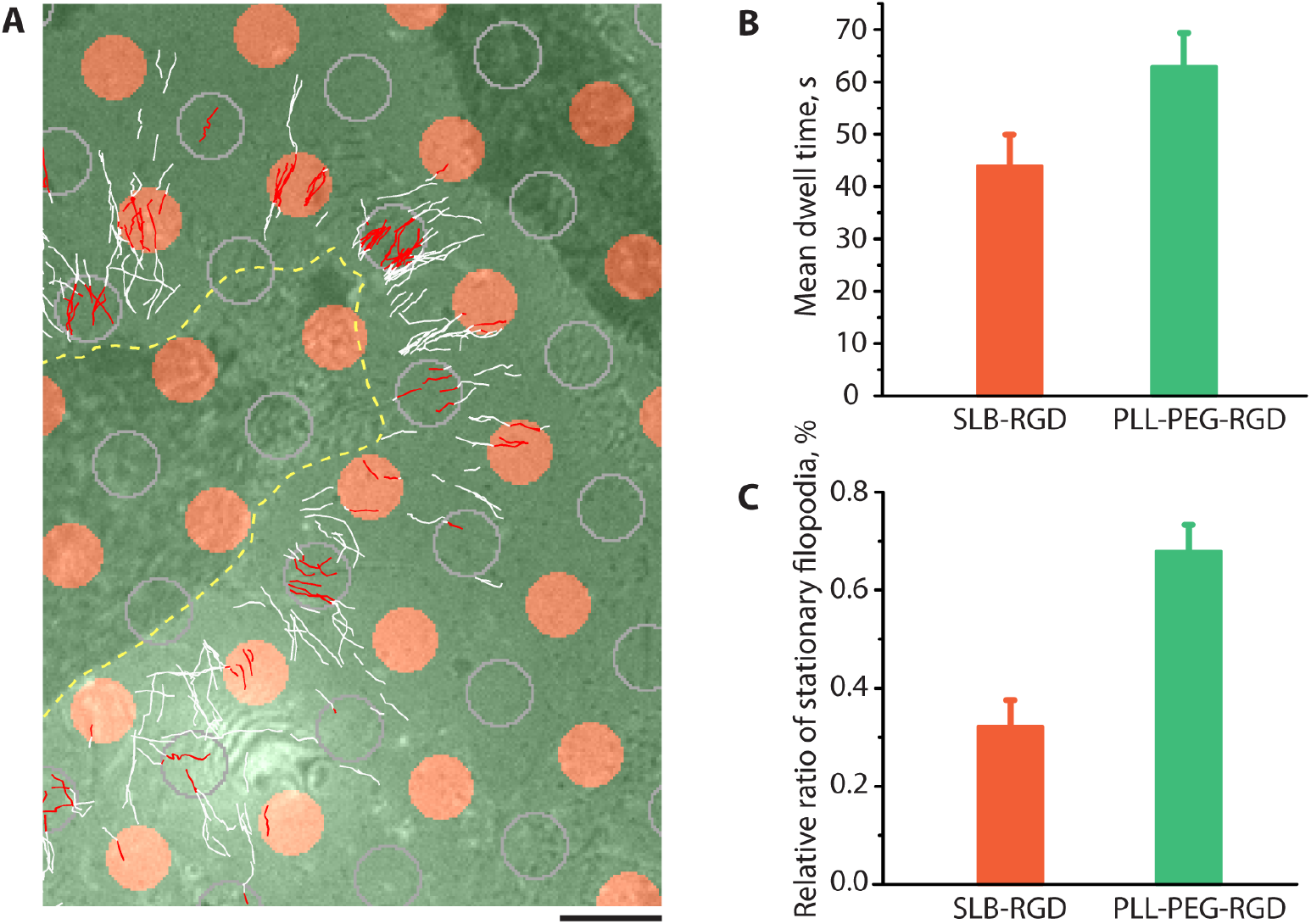
Attachment of filopodia to RGD-coated rigid and fluid substrate. **(A)** HeLa-JW cells expressing GFP-myosin X were plated on micropatterned coverslips covered with circular islands (D = 3μm) of supported lipid bilayer (SLB) conjugated to RGD (orange circles), organized into a square lattice. The glass between the islands was covered with poly-L-lysine-PEG conjugated to RGD at the same density (fig. S8). Trajectories of GFP-positive filopodia tips acquired during a 14-36 min time interval are shown. The cell border is shown by a yellow dashed line. For comparison of the trajectories on rigid and fluid substrates, the circles of similar diameter were drawn by computer in the centers of the square lattice formed by SLB islands (outlined by gray contours). The segments of the trajectories located inside either the SLB islands or the drawn circles on the rigid substrate are shown in red, and the remaining parts of the trajectories are shown in white. See also movie S14. Scale bar, 5μm. **(B-C)** Quantification of the trajectories of filopodia tips inside rigid and fluid circular islands for five cells (at least 200 individual trajectories per cell were scored). **(B)** The bars represent the average dwelling time that filopodia tips spent inside rigid (turquoise) or fluid (red) circles defined above. **(C)** Fraction of filopodia tip trajectories remaining inside rigid circles (green bar) and fluid circles (orange bar) relatively to the total number of trajectories in the circles during the period of observation. Error bars correspond to the SEM.

**Movie S1**

**Dynamics of filopodia induced by myosin X expression**

Growing, pausing and retracting filopodia in HeLa-JW cells overexpressing GFP-myosin X. The myosin X is localized to filopodia tips where it appears as characteristic “comet tails”. The duration of the entire movie is 3 min, the movie images were recorded at 4.31 frames per second (fps) and display rate is 50 fps.

**Movie S2**

**3D rendering of fixed myosin X-induced filopodia attached to a 2μ m fibronectin-coated bead**

HeLa-JW cell was transfected with GFP-myosin X and tdTomato-Ftractin. Single filopodium was attached to optically trapped bead. After pulling of the filopodium with the optical trap for several minutes, the cell was fixed by addition of 20% PFA (Tousimis) directly to the chamber. Image and position of the bead were reconstructed from bright field image. Note, clearly visible preserved integrity of actin core (red).

**Movie S3**

**Dynamics of force-induced growth of GFP-myosin X-induced filopodia**

Sustained filopodial growth was induced by pulling force generated by the optically trapped microbead coated with fibronectin (see details in SI and Fig. 1A). The frames from this movie, and the kymograph based on it, are shown in Fig. 1B and 1C respectively. The duration of the entire movie is 26 min 52 s. Movie images were recorded at 0.5 fps and displayed at 25 fps.

**Movie S4**

**Pulling of GFP-myosin X-induced filopodia using concanavalin A-coated beads**

Pulling force was applied to a filopodium of HeLa-JW cell via a cocanavalin-A-coated laser-trapped bead by moving the microscope stage. This force was applied to induce filopodia growth, but in contrast with the results obtained from experiments using fibronectin-coated beads (movie S3), under these conditions, filopodium did not grow and formation of membrane tethers occurred instead. In the movie, the tether formation can be inferred from the rapid return of the bead to the myosin X-containing tip of the filopodium after switching off the trap. Note, that the bead continued to be attached to the filopodium by thin membrane tether and stayed in the field of observation during last several seconds of the movie after the bead was released from the trap. The duration of the movie is 4 min and the movie images were recorded at 0.5 fps with a display rate of 15 fps.

**Movie S5**

**Pulling of active Cdc42 (Q61L)-induced filopodia**

Filopodia of a Hela-JW cell transfected with GFP-Cdc42 (Q61L) and tdTomato-Ftractin were pulled using the optical trap. Arrow indicates the center of the trap. The duration of the movie is 6 min 36 s and the movie images were recorded at 1 fps with a display rate of 15 fps.

**Movie S6**

**Centripetal co-movement of VASP and myosin X patches in cells treated with formin inhibitor**

Co-distribution of mApple-myosin X (green, left) and GFP-VASP (red, right) in the same filopodia of HeLa-JW cell 90 min after the addition of 20μM SMIFH2. The duration of the movie is 83 min 10 s and the movie images were recorded at 0.1 fps with a display rate of 50 fps. Movie was obtained by SDCM.

**Movie S7**

**Visualization of photoactivated actin in filopodia expressing myosin X**

This movie corresponds to the fig. S1C.

The left panel corresponds to GFP-myosin X. The right panel represents photoactivated PAmCherry-b-actin in the same filopodium of HeLa-JW cell. The site of photoactivation is indicated by the pink line and was performed at 6 s from the start of the movie. Note the retrograde movements of myosin X and actin inside the filopodium. The kymograph of the line drawn along the length of the filopodium, in which the photoactivation assay was performed, is shown in fig. S1C. The duration of the movie is 1 min 33 s and the movie images were recorded at 0.6 fps with a display rate of 7 fps.

**Movie S8A-G**

**Myosin II filaments in filopodia of HeLa-JW and Cos-7 cells**

These movies correspond to the Fig. 2A, C-H. (A) mApple-myosin X (red), RLC-GFP (green) and mTagBFP-Lifeact (blue) in HeLa-JW cell. (B-G) Cos-7 cells expressing myosin X (red) and myosin IIA or B (green) and their mutants: (B) GFP-myosin X. (C) mApple-myosin X and GFP-myosin IIA heavy chain. (D) GFP-myosin X ΔFERM. (E) GFP-myosin X ΔFERM and mCherry-myosin IIA heavy chain. (F) mApple-myosin X and GFP-myosin IIB heavy chain. (G) mApple-myosin X and GFP-myosin IIA N93K. Note the presence of myosin IIA filaments in movies A, C, E and G and the absence of myosin IIB in movie F at the filopodia bases. The duration of the movies were 15 min 50 s, 4 min 8s, 4 min 5 s, 4 min 5 s, 4 min 5s, 3 min 15 s and 4 min 5s respectively, the movie images were recorded and displayed at 0.003 and 7 for (A), 0.5 and 18 for (B), 0.2 and 7.5 fps for (C-G).

**Movies S9A-D**

**Inhibition of myosin II or formin suppresses filopodia adhesion and growth (see also Fig. 3 A-D)**

(A) Movie showing sustained growth of a filopodium induced by pulling force in a control HeLa-JW cell transfected with GFP-myosin X construct. The experiment was analogous to that described in the legend to Fig. 1A. The cell was used as a control for experiments with inhibitors presented in Fig. 3. The duration of the movie is 17 min 8 s. (B) Effect of light-insensitive blebbistatin. The cell was pretreated with 20μM S-nitro-blebbistatin for 10-20 min prior to placing the laser trapped fibronectin-coated bead onto the filopodium tip. 30 s later, stage movement commenced simultaneously with filming. Note that adhesion of the bead to filopodium was broken 6 min 45 s after starting the stage movement. The duration of the movie is 13 min 12 s. (C) Effect of myosin IIA knockdown. The detachment of the filopodium from the bead occurred 3 min 50 s after the stage movement was initiated. The velocity of the movement of myosin X patches observed in this experiment was significantly lower then the velocity of myosin II-driven movement (Fig. 4). The duration of the movie is 4 min 38 s. (D) Effect of formin inhibition by SMIFH2. The cell was pretreated with 40μM SMIFH2 for 1 hour prior the bead being placed onto the filopodium tip. To check whether the bead had attached to the filopodium, the laser trap was switched off for several seconds, during which time unattached beads typically disappeared from the field of view. The detachment of the filopodium from the bead occurred about 2 min after the stage movement was initiated. The duration of the entire movie is 4 min 56 s. All the movies in this figure were recorded at 0.5 fps with a display rate of 15 fps.

**Movie S10**

**Effect of ROCK inhibitor, Y27632, on force-induced filopodia growth and adhesion**

This movie corresponds to the frames shown in fig. S4. HeLa-JW cell filopodium growth was induced by applying a pulling force, generated as a result of microscope stage movement, as in Fig. 1. At about 4 min after stage movement commenced, 50μM of Y27632 was added, which eventually resulted in bead detachment at 17 min. The intensity of actin labeling was relatively low and apparent “disappearance” of actin in the second half of the movie was a result of photobleacning. The duration of the entire movie is 19 min 22 s. The movie was recorded at 1 fps and displayed at 25 fps.

**Movie S11**

**Effect of formin inhibitor SMIFH2 on force-induce filopodia growth and adhesion**

40uM of SMIFH2 was added to HeLa-JW cell overexpressing GFP-myosin X (green) and tdTomato-Fractin (red) at 11 min of the movie after establishment of sustained filopodia growth as in fig. S4. Several minutes after, cessation of growth of the actin core and the drop in the pulling force generated by the filopodium (data not shown) were prominent but the fibronectin-coated bead remained associated with the filopodium tip via a membrane tether. The duration of the entire movie is 20 min 39 s. The movie was recorded at 1 fps and displayed at 30 fps.

**Movie S12**

**Effect of formin inhibition on dynamics of myosin X in filopodia**

This movie corresponds to the Fig. 4B. The filopodia were induced by HeLa-JW cell transfection with a construct encoding GFP-myosin X. Before imaging, the cell was treated with 20μM of SMIFH2 for 2 hours. Note the apparent disintegration of the myosin X containing patch at the tip of the filopodium into numerous small patches, the majority of which rapidly moved centripetally from the filopodium tip to the cell body, with retrograde movement of myosin X patches sometimes interrupted by short periods of anterograde sliding towards the filopodium tip. The duration of the entire movie is 5 min; the movie was recorded at 4 fps and displayed at 100 fps. The movie was obtained with SDCM.

**Movie S13A-C**

**Effect of myosin II inhibition on fast centripetal movement of myosin X patches in filopodia of SMIFH2 treated cells**

These movies correspond to Fig.4C. A HeLa-JW cell transfected with GFP-myosin X was filmed before treatment (A), 15 min after the addition of 20μM SMIFH2 (B) and 20 min after the subsequent addition of 30μM Y27632 (C). Note that after SMIFH2 was added, myosin X comet tails underwent rapid disintegration into small patches, which moved centripetally towards the cell body (B). Addition of Y27632 resulted in cessation of this movement (C). The duration of the movies A-C were 6 min 41 s; the movie images were recorded at 0.1 fps and displayed at 7 fps. The movies were obtained with SDCM.

**Movie S14**

**Filopodia distinguish between fluid and rigid substrates**

GFP-myosin X transfected HeLa-JW cell spreading on the micropatterned substrate with 3um islands covered with supported lipid bilayer (SLB). Both the islands and the rigid substrate between them were coated with fluorescent Dylight-405 RGD ligand shown in blue. The GFP-myosin X positive filopodia tips are shown in green. The movie started 30 min following the cell plating. Note that filopodia tips apparently avoid the SLB islands, see the analysis in fig. S9. The duration of the movies was 16 min 40 s; the movie images were recorded at 0.3 fps and displayed at 30 fps.

## REFERENCES

1. Jacquemet, G., Hamidi, H. & Ivaska, J. Filopodia in cell adhesion, 3D migration and cancer cell invasion. Curr Opin Cell Biol 36, 23–31 (2015).

2. Mattila, P.K. & Lappalainen, P. Filopodia: molecular architecture and cellular functions. Nat Rev Mol Cell Biol 9, 446–454 (2008).

3. Heckman, C.A. & Plummer, H.K., 3rd Filopodia as sensors. Cell Signal 25, 2298–2311 (2013).

4. Bryant, P.J. Filopodia: fickle fingers of cell fate? Curr Biol 9, R655–657 (1999).

5. Prols, F., Sagar & Scaal, M. Signaling filopodia in vertebrate embryonic development. Cell Mol Life Sci 73, 961–974 (2016).

6. Romero, S. et al. Filopodium retraction is controlled by adhesion to its tip. J Cell Sci 125, 4999–5004 (2012).

7. Vonna, L., Wiedemann, A., Aepfelbacher, M. & Sackmann, E. Micromechanics of filopodia mediated capture of pathogens by macrophages. Eur Biophys J 36, 145–151 (2007).

8. Moller, J., Luhmann, T., Chabria, M., Hall, H. & Vogel, V. Macrophages lift off surface-bound bacteria using a filopodium-lamellipodium hook-and-shovel mechanism. Sci Rep 3, 2884 (2013).

9. Bornschlogl, T. et al. Filopodial retraction force is generated by cortical actin dynamics and controlled by reversible tethering at the tip. Proc Natl Acad Sci U S A 110, 18928–18933 (2013).

10. Faix, J., Breitsprecher, D., Stradal, T.E. & Rottner, K. Filopodia: Complex models for simple rods. Int J Biochem Cell Biol 41, 1656–1664 (2009).

11. Jaiswal, R. et al. The formin Daam1 and fascin directly collaborate to promote filopodia formation. Curr Biol 23, 1373–1379 (2013).

12. Vignjevic, D. et al. Role of fascin in filopodial protrusion. J Cell Biol 174, 863–875 (2006).

13. Delanote, V. et al. An alpaca single-domain antibody blocks filopodia formation by obstructing L-plastin-mediated F-actin bundling. FASEB J 24, 105–118 (2010).

14. Van Audenhove, I. et al. Fascin Rigidity and L-plastin Flexibility Cooperate in Cancer Cell Invadopodia and Filopodia. J Biol Chem 291, 9148–9160 (2016).

15. Barzik, M., McClain, L.M., Gupton, S.L. & Gertler, F.B. Ena/VASP regulates mDia2-initiated filopodial length, dynamics, and function. Mol Biol Cell 25, 2604–2619 (2014).

16. Block, J. et al. Filopodia formation induced by active mDia2/Drf3. J Microsc 231, 506–517 (2008).

17. Pellegrin, S. & Mellor, H. The Rho family GTPase Rif induces filopodia through mDia2. Curr Biol 15, 129–133 (2005).

18. Schirenbeck, A., Bretschneider, T., Arasada, R., Schleicher, M. & Faix, J. The Diaphanous-related formin dDia2 is required for the formation and maintenance of filopodia. Nat Cell Biol 7, 619–625 (2005).

19. Block, J. et al. FMNL2 drives actin-based protrusion and migration downstream of Cdc42. Curr Biol 22, 1005–1012 (2012).

20. Harris, E.S., Gauvin, T.J., Heimsath, E.G. & Higgs, H.N. Assembly of filopodia by the formin FRL2 (FMNL3). Cytoskeleton (Hoboken) 67, 755–772 (2010).

21. Young, L.E., Heimsath, E.G. & Higgs, H.N. Cell type-dependent mechanisms for formin-mediated assembly of filopodia. Mol Biol Cell 26, 4646–4659 (2015).

22. Applewhite, D.A. et al. Ena/VASP proteins have an anti-capping independent function in filopodia formation. Mol Biol Cell 18, 2579–2591 (2007).

23. Schirenbeck, A., Arasada, R., Bretschneider, T., Schleicher, M. & Faix, J. Formins and VASPs may co-operate in the formation of filopodia. Biochem Soc Trans 33, 1256–1259 (2005).

24. Bear, J.E. & Gertler, F.B. Ena/VASP: towards resolving a pointed controversy at the barbed end. J Cell Sci 122, 1947–1953 (2009).

25. Zhang, H. et al. Myosin-X provides a motor-based link between integrins and the cytoskeleton. Nat Cell Biol 6, 523–531 (2004).

26. Kerber, M.L. & Cheney, R.E. Myosin-X: a MyTH-FERM myosin at the tips of filopodia. J Cell Sci 124, 3733–3741 (2011).

27. Sousa, A.D. & Cheney, R.E. Myosin-X: a molecular motor at the cell's fingertips. Trends Cell Biol 15, 533–539 (2005).

28. Hoffmann, B. & Schafer, C. Filopodial focal complexes direct adhesion and force generation towards filopodia outgrowth. Cell Adh Migr 4, 190–193 (2010).

29. Sydor, A.M., Su, A.L., Wang, F.S., Xu, A. & Jay, D.G. Talin and vinculin play distinct roles in filopodial motility in the neuronal growth cone. J Cell Biol 134, 1197–1207 (1996).

30. Lagarrigue, F. et al. A RIAM/lamellipodin-talin-integrin complex forms the tip of sticky fingers that guide cell migration. Nat Commun 6, 8492 (2015).

31. Schafer, C. et al. One step ahead: role of filopodia in adhesion formation during cell migration of keratinocytes. Exp Cell Res 315, 1212–1224 (2009).

32. Schafer, C. et al. The key feature for early migratory processes: Dependence of adhesion, actin bundles, force generation and transmission on filopodia. Cell Adh Migr 4, 215–225 (2010).

33. Geiger, B., Spatz, J.P. & Bershadsky, A.D. Environmental sensing through focal adhesions. Nat Rev Mol Cell Biol 10, 21–33 (2009).

34. Sun, Z., Guo, S.S. & Fassler, R. Integrin-mediated mechanotransduction. J Cell Biol 215, 445–456 (2016).

35. Riveline, D. et al. Focal contacts as mechanosensors: externally applied local mechanical force induces growth of focal contacts by an mDia1-dependent and ROCK-independent mechanism. J Cell Biol 153, 1175–1186 (2001).

36. Goffin, J.M. et al. Focal adhesion size controls tension-dependent recruitment of alpha-smooth muscle actin to stress fibers. J Cell Biol 172, 259–268 (2006).

37. Prager-Khoutorsky, M. et al. Fibroblast polarization is a matrix-rigidity-dependent process controlled by focal adhesion mechanosensing. Nat Cell Biol 13, 1457–1465 (2011).

38. Trichet, L. et al. Evidence of a large-scale mechanosensing mechanism for cellular adaptation to substrate stiffness. Proc Natl Acad Sci U S A 109, 6933–6938 (2012).

39. Wong, S., Guo, W.H. & Wang, Y.L. Fibroblasts probe substrate rigidity with filopodia extensions before occupying an area. Proc Natl Acad Sci U S A 111, 17176–17181 (2014).

40. Berg, J.S. & Cheney, R.E. Myosin-X is an unconventional myosin that undergoes intrafilopodial motility. Nat Cell Biol 4, 246–250 (2002).

41. Watanabe, T.M., Tokuo, H., Gonda, K., Higuchi, H. & Ikebe, M. Myosin-X induces filopodia by multiple elongation mechanism. J Biol Chem 285, 19605–19614 (2010).

42. Tullio, A.N. et al. Nonmuscle myosin II-B is required for normal development of the mouse heart. Proc Natl Acad Sci U S A 94, 12407–12412 (1997).

43. Even-Ram, S. et al. Myosin IIA regulates cell motility and actomyosin-microtubule crosstalk. Nat Cell Biol 9, 299–309 (2007).

44. Bohil, A.B., Robertson, B.W. & Cheney, R.E. Myosin-X is a molecular motor that functions in filopodia formation. Proc Natl Acad Sci U S A 103, 12411–12416 (2006).

45. Choi, C.K. et al. Actin and alpha-actinin orchestrate the assembly and maturation of nascent adhesions in a myosin II motor-independent manner. Nat Cell Biol 10, 1039–1050 (2008).

46. Vicente-Manzanares, M., Zareno, J., Whitmore, L., Choi, C.K. & Horwitz, A.F. Regulation of protrusion, adhesion dynamics, and polarity by myosins IIA and IIB in migrating cells. J Cell Biol 176, 573–580 (2007).

47. Smith, R.C. et al. Regulation of myosin filament assembly by light-chain phosphorylation. Philos Trans R Soc Lond B Biol Sci 302, 73–82 (1983).

48. Betapudi, V. Life without double-headed non-muscle myosin II motor proteins. Front Chem 2, 45 (2014).

49. Vicente-Manzanares, M., Ma, X., Adelstein, R.S. & Horwitz, A.R. Non-muscle myosin II takes centre stage in cell adhesion and migration. Nat Rev Mol Cell Biol 10, 778–790 (2009).

50. Hu, S. et al. Long-range self-organization of cytoskeletal myosin II filament stacks. Nat Cell Biol 19, 133–141 (2017).

51. Rizvi, S.A. et al. Identification and characterization of a small molecule inhibitor of formin-mediated actin assembly. Chem Biol 16, 1158–1168 (2009).

52. Tokuo, H. & Ikebe, M. Myosin X transports Mena/VASP to the tip of filopodia. Biochem Biophys Res Commun 319, 214–220 (2004).

53. Yu, C.H., Law, J.B., Suryana, M., Low, H.Y. & Sheetz, M.P. Early integrin binding to Arg-Gly-Asp peptide activates actin polymerization and. Proc Natl Acad Sci U S A 108, 20585–20590 (2011).

54. Jacquemet, G. et al. L-type calcium channels regulate filopodia stability and cancer cell invasion downstream of integrin signalling. Nat Commun 7, 13297 (2016).

55. Billington, N., Wang, A., Mao, J., Adelstein, R.S. & Sellers, J.R. Characterization of three full-length human nonmuscle myosin II paralogs. J Biol Chem 288, 33398–33410 (2013).

56. Hundt, N., Steffen, W., Pathan-Chhatbar, S., Taft, M.H. & Manstein, D.J. Load-dependent modulation of non-muscle myosin-2A function by tropomyosin 4.2. Sci Rep 6, 20554 (2016).

57. Kovacs, M., Wang, F., Hu, A., Zhang, Y. & Sellers, J.R. Functional divergence of human cytoplasmic myosin II: kinetic characterization of the non-muscle IIA isoform. J Biol Chem 278, 38132–38140 (2003).

58. Yan, J., Yao, M., Goult, B.T. & Sheetz, M.P. Talin Dependent Mechanosensitivity of Cell Focal Adhesions. Cell Mol Bioeng 8, 151–159 (2015).

59. Atherton, P. et al. Vinculin controls talin engagement with the actomyosin machinery. Nat Commun 6, 10038 (2015).

60. Hu, X. et al. Cooperative Vinculin Binding to Talin Mapped by Time-Resolved Super Resolution Microscopy. Nano Lett 16, 4062–4068 (2016).

61. Gauthier, N.C. & Roca-Cusachs, P. Mechanosensing at integrin-mediated cell-matrix adhesions: from molecular to integrated mechanisms. Curr Opin Cell Biol 50, 20–26 (2018).

62. Hu, W., Wehrle-Haller, B. & Vogel, V. Maturation of filopodia shaft adhesions is upregulated by local cycles of. PLoS One 9, 0107097 (2014).

63. Goult, B.T. et al. RIAM and vinculin binding to talin are mutually exclusive and regulate adhesion. J Biol Chem 288, 8238–8249 (2013).

64. Lafuente, E.M. et al. RIAM, an Ena/VASP and Profilin ligand, interacts with Rap1-GTP and mediates Rap1-induced adhesion. Dev Cell 7, 585–595 (2004).

65. Courtemanche, N., Lee, J.Y., Pollard, T.D. & Greene, E.C. Tension modulates actin filament polymerization mediated by formin and profilin. Proc Natl Acad Sci U S A 110, 9752–9757 (2013).

66. Jegou, A., Carlier, M.F. & Romet-Lemonne, G. Formin mDia1 senses and generates mechanical forces on actin filaments. Nat Commun 4, 1883 (2013).

67. Kozlov, M.M. & Bershadsky, A.D. Processive capping by formin suggests a force-driven mechanism of actin polymerization. J Cell Biol 167, 1011–1017 (2004).

68. Yu, M. et al. mDia1 senses both force and torque during F-actin filament polymerization. Nat Commun 8, 1650 (2017).

69. Galbraith, C.G., Yamada, K.M. & Galbraith, J.A. Polymerizing actin fibers position integrins primed to probe for adhesion sites. Science 315, 992–995 (2007).

70. Guillou, H. et al. Lamellipodia nucleation by filopodia depends on integrin occupancy and downstream Rac1 signaling. Exp Cell Res 314, 478–488 (2008).

71. Chan, C.E. & Odde, D.J. Traction dynamics of filopodia on compliant substrates. Science 322, 1687–1691 (2008).

72. Lo, C.M., Wang, H.B., Dembo, M. & Wang, Y.L. Cell movement is guided by the rigidity of the substrate. Biophys J 79, 144–152 (2000).

73. Heidemann, S.R. & Bray, D. Tension-driven axon assembly: a possible mechanism. Front Cell Neurosci 9, 316 (2015).

74. Kerstein, P.C., Nichol, R.I. & Gomez, T.M. Mechanochemical regulation of growth cone motility. Front Cell Neurosci 9, 244 (2015).

75. Bridgman, P.C., Dave, S., Asnes, C.F., Tullio, A.N. & Adelstein, R.S. Myosin IIB is required for growth cone motility. J Neurosci 21, 6159–6169 (2001).

76. Bray, D. Mechanical tension produced by nerve cells in tissue culture. J Cell Sci 37, 391–410 (1979).

77. Bai, M., Harfe, B. & Freimuth, P. Mutations that alter an Arg-Gly-Asp (RGD) sequence in the adenovirus type 2 penton base protein abolish its cell-rounding activity and delay virus reproduction in flat cells. J Virol 67, 5198–5205 (1993).

78. Paran, Y. et al. Development and application of automatic high-resolution light microscopy for cell-based screens. Methods Enzymol 414, 228–247 (2006).

79. Schell, M.J., Erneux, C. & Irvine, R.F. Inositol 1,4,5-trisphosphate 3-kinase A associates with F-actin and dendritic spines via its N terminus. J Biol Chem 276, 37537–37546 (2001).

80. Lin, A.Y., Prochniewicz, E., James, Z.M., Svensson, B. & Thomas, D.D. Large-scale opening of utrophin's tandem calponin homology (CH) domains upon actin binding by an induced-fit mechanism. Proc Natl Acad Sci U S A 108, 12729–12733 (2011).

81. Winder, S.J. et al. Calmodulin regulation of utrophin actin binding. Biochem Soc Trans 23, 397S (1995).

82. Kengyel, A., Wolf, W.A., Chisholm, R.L. & Sellers, J.R. Nonmuscle myosin IIA with a GFP fused to the N-terminus of the regulatory light chain is regulated normally. J Muscle Res Cell Motil 31, 163–170 (2010).

83. Watanabe, S. et al. mDia2 induces the actin scaffold for the contractile ring and stabilizes its position during cytokinesis in NIH 3T3 cells. Mol Biol Cell 19, 2328–2338 (2008).

84. Brock, R. & Jovin, T.M. Heterogeneity of signal transduction at the subcellular level: microsphere-based focal EGF receptor activation and stimulation of Shc translocation. J Cell Sci 114, 2437–2447 (2001).

85. Neuman, K.C. & Block, S.M. Optical trapping. Rev Sci Instrum 75, 2787–2809 (2004).

86. Toth, M.A. et al. Biochemical Activities of the Wiskott-Aldrich Syndrome Homology Region 2 Domains of Sarcomere Length Short (SALS) Protein. J Biol Chem 291, 667–680 (2016).

87. Spudich, J.A. & Watt, S. The regulation of rabbit skeletal muscle contraction. I. Biochemical studies of the interaction of the tropomyosin-troponin complex with actin and the proteolytic fragments of myosin. J Biol Chem 246, 4866–4871 (1971).

88. Romero, S. et al. Formin is a processive motor that requires profilin to accelerate actin assembly and associated ATP hydrolysis. Cell 119, 419–429 (2004).

89. Tsygankov, D. et al. CellGeo: a computational platform for the analysis of shape changes in cells with complex geometries. J Cell Biol 204, 443–460 (2014).

90. Lin, W.C., Yu, C.H., Triffo, S. & Groves, J.T. Supported membrane formation, characterization, functionalization, and patterning. Curr Protoc Chem Biol 2, 235–269 (2010).

91. Gelles, J., Schnapp, B.J. & Sheetz, M.P. Tracking kinesin-driven movements with nanometre-scale precision. Nature 331, 450–453 (1988).

